# The Genomic Landscape of Post-Black Death Epidemics in Northern Europe and the Caucasus

**DOI:** 10.64898/2026.08.25.746822

**Authors:** Cameron Robert Ferguson, Åshild J. Vågene, Marcel Keller, Caroline Ahlström Arcini, Eveline Altena, Gorka Lasso, Axel Christophersen, Maria Cinthio, Sean Denham, Sascha Dreyer Nielsen, S. Sunna Ebenesersdóttir, Martin R. Ellegaard, Shyam Gopalakrishnan, Agnar Helgason, Miren Iraeta-Orbegozo, Liam Thomas Lanigan, Xiaodong Liu, Emma Loftus, Maja M. B. Lunn, Ashot Margaryan, Kristjan Moore, Bente Philippsen, Hannes Schroeder, Hakob Simonyan, Zaruhi Khachatryan, Pooja Swali, Cedric CS Tan, Levon Yepiskoposyan, Laurent Frantz, Philip Slavin, Francois Balloux, M Thomas P Gilbert, Lucy van Dorp

**Author notes:** Contributed equally (Cameron Robert Ferguson, Åshild J. Vågene). also contributed equally to this work (Francois Balloux, M Thomas P Gilbert, Lucy van Dorp).

## Abstract

One of the most devastating events in human history, the Black Death (c.1347-1353), marked the beginning of the Second Plague Pandemic. After initially receding, plague returned in intermittent outbreaks throughout Europe, beginning with the *pestis secunda*. Despite its significance, the post-Black Death epidemiology of *Yersinia pestis* remains poorly understood. Here, we report 23 new *Y. pestis* genomes recovered from Second Plague Pandemic contexts across Scandinavia, the Netherlands, Iceland, and Armenia, spanning approximately 270 years. Applying a reproducible mutation-filtering pipeline to assess genetic diversity, we report both Black Death and post-Black Death lineages and document multiple waves of plague at individual cemetery sites. We identify four *pestis secunda* genomes, including three from Armenia, supporting the eastward dissemination of this lineage prior to its disappearance from Europe. We also resolve a previously under-characterised Branch 1A sub-lineage of *Y. pestis* and provide the first genomic evidence of plague in Iceland, resolving longstanding uncertainty over its presence on the island. Finally, by calling variants against a reconstructed ancestral reference, we identify clade-defining mutations, including nonsynonymous changes in metabolic and biofilm-related genes with predicted structural effects that may have contributed to shaping the epidemiological dynamics of the Second Plague Pandemic.

## Introduction

*Yersinia pestis*, the causative agent of plague, is a zoonotic pathogen maintained in enzootic cycles among wild rodent populations and their flea vectors, with periodic spillover into humans^1,2^. In humans, plague can manifest in several clinical forms, including bubonic, pneumonic, and septicemic plague. Without treatment, mortality rates range from 30% to nearly 100%, with pneumonic plague typically being the most lethal^3^. Historical, archaeological and genomic evidence indicates that large-scale epidemic expansions have occurred intermittently over the course of its evolutionary history, with at least three well-characterised plague pandemics to date; the First Plague Pandemic (Europe and wider Mediterranean region, 6^th^ to 8^th^ centuries CE)^4,5^, the Second Plague Pandemic (Europe, mid-14^th^ century Black Death until the early 19^th^ century CE)^6–8^ and the Third Plague Pandemic which spread worldwide between the late 18th and early 20th centuries^9–11^. As a consequence, *Y. pestis* was established in novel endemic foci spanning Africa, Asia and the Americas where it continues to pose a significant challenge to humans^10,12,13^. Based on observations and experiments during the Third Plague Pandemic, the classical model associates human plague epidemics with black rats (*Rattus rattus*) as a commensal rodent reservoir and their fleas (*Xenopsylla cheopis*) as vectors^14^.

Among the pandemics attributed to *Y. pestis*, the European Black Death (1347-1353) stands out as one of the most devastating demographic events in human history, resulting in the loss of up to 60% of the population in some regions^8^. It is considered the fulminant start of the Second Plague Pandemic which went on to affect large parts of Africa and Eurasia from the 14th to the early 19th centuries. While historical sources provide detailed accounts of the scale and progression of these outbreaks and their consequences^8,15,16^, analyses of ancient pathogen DNA recovered from archaeological remains offer direct insight into their epidemiology and transmission dynamics^17–22^.

The Black Death is thought to have originated in Central Asia^23–25^. Genomic analyses of *Y. pestis* recovered from Kara-Djigach, Kyrgyzstan, confidently dated to 1338, identified strains directly ancestral to those responsible for the European pandemic^19^. These findings are consistent with a model of westward spread of the Black Death along established trade routes, with early outbreaks recorded in Messina, Sicily. Following importation from Caffa in 1347^8,26,27^, a single clonal lineage spread rapidly across Europe from 1348 to 1353^8,21,22^.

Historical sources document a rapid resurgence of plague within a decade of the Black Death, the first of which was known as the *pestis secunda,* with major outbreaks recorded between 1356 and 1366^28,29^. Genomic analyses of infected individuals from Krakauer Berg, Germany, suggested a Central European origin for the *pestis secunda*^18^, supporting historical assessments^28^ but contrasting with earlier hypotheses proposing repeated reintroductions from Asian sources^30–33^. Although its precise origin remains debated, genomic evidence suggests that the *pestis secunda* then spread rapidly, with *Y. pestis* genomes from the *pestis secunda* found in remains from multiple locations, including Bergen op Zoom in the Netherlands^33^, London in England^17^, and Bolgar in western Russia^22^. These *pestis secunda* strains are derived from the Black Death genotype, but do not fall within the diversity of genomes obtained from later Second Plague Pandemic outbreaks in Europe. Instead, they form their own lineage differing from the Black Death at two diagnostic SNPs, and occupying a phylogenetic position ancestral to Branch 1B, including the Third Plague Pandemic lineages^17,18,22,33^. Historical accounts suggest that the *pestis secunda* may have differed epidemiologically from the Black Death, with some studies indicating disproportionate mortality among younger individuals and males^29^. The extent to which these patterns reflect differences in exposure, acquired immunity, or underlying pathogen biology remains unclear.

Regarding the origins of the Third Plague Pandemic, historical records and genomic analyses suggest that it emerged in the late 18th century in a region near present day Yunnan Province, China, before spreading globally via expanding international trade networks in the mid-19th century^11,12^. Phylogenetically, the Third Plague Pandemic lineage descends from a branch associated with the *pestis secunda*, raising questions about where this lineage persisted between its apparent disappearance in Europe and its re-emergence in East Asia centuries later^34^.

Despite growing knowledge on the early diversification of *Y. pestis* during the Second Plague Pandemic^17–22^, most post-Black Death outbreak-associated clusters remain unevenly distributed. To date, approximately 70 high coverage (>5x) ancient *Y. pestis* genomes are available from post-Black Death contexts, representing 38 archaeological sites across 13 countries. While this constitutes a substantial genomic dataset, sampling remains geographically restricted and is particularly sparse at the margins of medieval Europe, including its northern and eastern peripheries, such as the North Atlantic islands and into the Caucasus. These regions were likely important for the persistence, dispersal and reintroduction of plague^35–37^. This is particularly true for Iceland, where the historically documented occurrence of plague has fuelled theories about alternative transmission cycles due to the apparent lack of rats and even some speculation as to whether these outbreaks were indeed caused by *Y. pestis*^38,39^.

Here, we present 23 *Y. pestis* genomes from five sites across Europe and the Caucasus, including longitudinal sampling spanning approximately 270 years (1270-1536) from Trondheim (Norway) and Lund (former Denmark, present-day Sweden), both of which experienced substantial population declines during the Black Death and subsequent outbreaks^35,36,40–43^. Additionally, we provide the first genomic evidence confirming the presence of post-Black Death plague in Iceland, as well as new representatives of the *pestis secunda* from Sweden and Armenia. In total, we report 14 new ancient genomes with >5x coverage and another nine with lower coverage but that could still be confidently placed in the *Y. pestis* phylogenetic tree. Using a novel Ancestral Reference Mapping (ARM) framework, rather than standard allele calling against a modern reference genome, we characterise the mutations defining major clades over the *Y. pestis* phylogeny. This further allows us to pinpoint specific non-synonymous changes arising during the Black Death and the *pestis secunda*, which are shared by all Third Plague Pandemic strains in circulation today.

## Results

### Recovery of *Y. pestis* genomes from the Black Death and post-Black Death era

We initially screened DNA sequence data generated from several recently published and ongoing ancient DNA analyses of human remains from Europe^44^ and the South Caucasus. These included individuals from Trondheim (Norway)^45,46^, Trinitatis Church in Lund (Sweden)^47^, Aghtsk in Armenia^48,49^, St. Catharina’s Church in Eindhoven (the Netherlands)^50^, and the cemetery at Útskála in Útskálakirkjugarður (Iceland)^51^. Shotgun sequenced metagenomic DNA from all individuals was screened for the presence of *Y. pestis* reads using the MALT/HOPS pipeline^52,53^ to assign reads to microbial taxonomic nodes of best fit, authenticating them using established criteria, including elevated *Y. pestis* and *Y. pseudotuberculosis complex* read counts, characteristic edit-distance distributions, and damage patterns consistent with ancient DNA. Based on these results we identified a set of candidate *Y. pestis* positive individuals for targeted capture enrichment (see **Methods**, **Supplementary Note 1-2**, and **Supplementary Tables S1-3**).

Following capture enrichment (see **Methods, Supplementary Note 2**), we recovered 23 *Y. pestis* genomes, all with at least 1x mean genome-wide coverage, of which 15 exceeded 3x (**Supplementary Tables S1-2, Figure 1**). These included 14 genomes from Trinitatis Church in Lund (Sweden), of which 12 exceeded 3x mean coverage, spanning the period c. 1270-1536 and enabling comparison of strains across the end of the High and Late Middle Ages, extending into the Early Modern Period^54^. From Trondheim (Norway), we identified four *Y. pestis* infections (two >30x and two ∼1x coverage) dated, based on radiocarbon and archaeological context, to between 1275-1600. We further recovered three genomes from Aghtsk c.1260-1365 (Armenia) at approximately 1-5x coverage representing the first second plague pandemic genomes from Armenia, one genome from the Catharina cemetery site in Eindhoven (∼2.5x), and the first direct molecular evidence of *Y. pestis* in Iceland (Útskála, ∼2.5x), radiocarbon dated to 1460-1814 calAD following marine reservoir correction.

**Figure 1:**
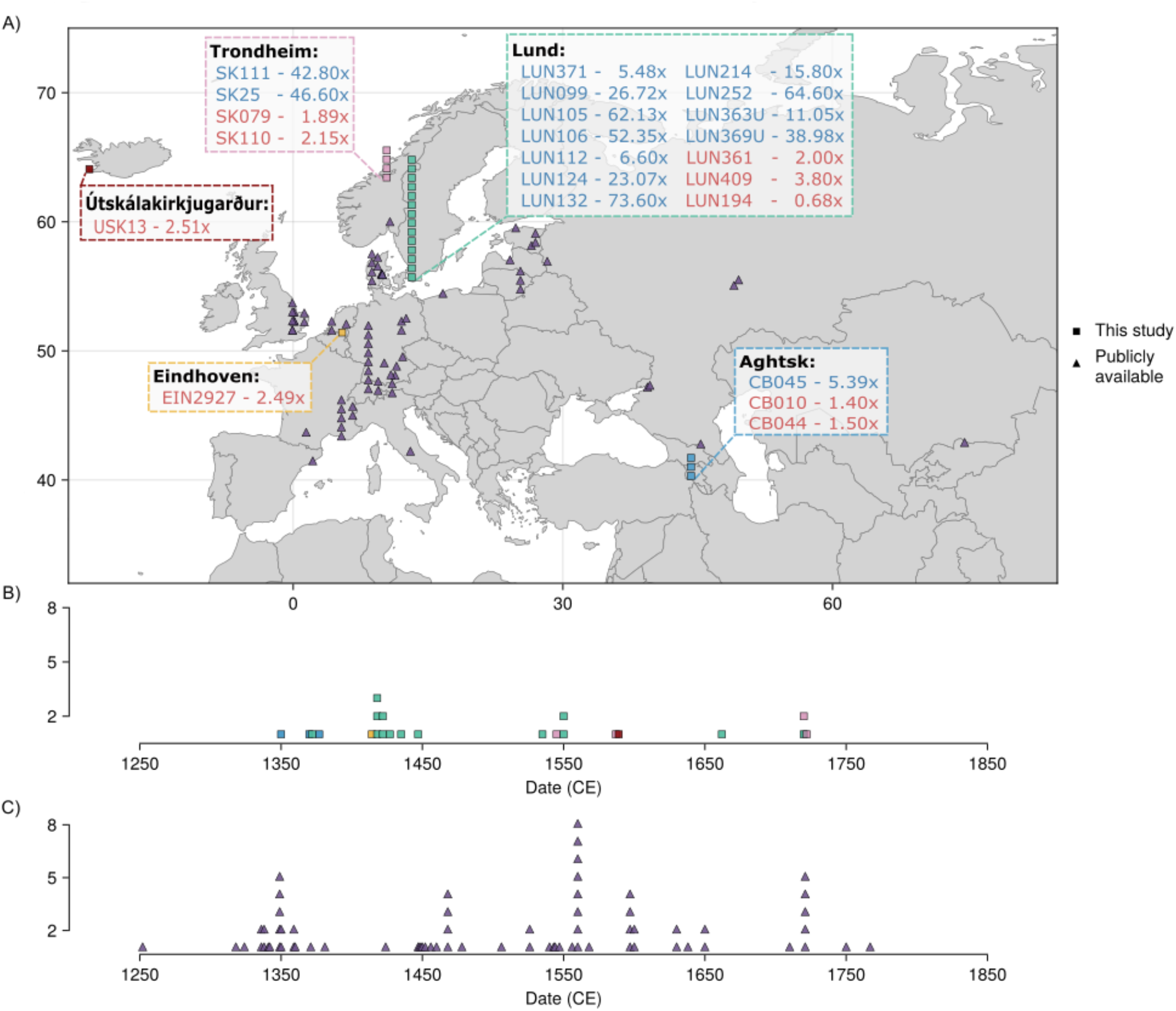
The spatio-temporal distribution of Second Plague Pandemic genomes. A) Map providing the sampling locations for *Y. pestis* genomes recovered in this study. Samples from the same site are stacked vertically and coloured according to the site. Publicly available ancient *Y. pestis* genomes included are shown as purple triangles. The names of genomes recovered in this study are provided in text boxes linking to the sites considered and are coloured blue if they are above 5x and red if they are below 5x. Squares indicate novel samples from this study. B) Timeline depicting the distribution of Second Plague Pandemic *Y. pestis* genomes recovered in this study across time, based on the midpoint of their radiocarbon estimates or archaeological date range. Samples with the same sampling dates are stacked vertically, and the points are coloured according to the site where they were recovered. C) Timeline providing the distribution of publicly available Second Plague Pandemic genomes across time, based on the midpoint of their radiocarbon estimates or archaeological date range, from published literature. Samples with the same sampling dates are stacked vertically. A full description of *Y. pestis* genomes considered is available in Supplementary Table 1.

All libraries were mapped to the *Y. pestis* CO92 reference genome (GCF_000009065.1) and exhibited even coverage profiles, appropriate edit-distance distributions, and post-mortem damage patterns consistent with authentic ancient DNA (see **Methods**, **Supplementary Table S1, Supplementary Notes 2-3**). Together, these data substantially expand the available genomic record of *Y. pestis* during the Second Plague Pandemic, particularly through the inclusion of previously unreported regions such as Iceland and Armenia. Notably, the *Y. pestis* observations recovered from Lund provide a rare longitudinal series over the Second Plague Pandemic of 12 genomes at >3x coverage spanning approximately 270 years.

### Phylogenetic reconstruction of Black Death and post-Black Death epidemics

We assessed our new *Y. pestis* observations in the context of the genomic diversity of previously recovered and reported genomic data from the Black Death and post-Black Death periods, further contextualised with 204 modern *Y. pestis* strains (**Supplementary Tables S4-5**).

Ancient metagenomic data, particularly at low genome-wide coverage, present challenges in confidently identifying true variants in *Y. pestis*. We used several approaches to ensure the robustness of the alignment prior to ascertaining phylogeographic patterns. In particular, a consistent and reproducible filtering framework was applied (see **Methods, Supplementary Note 3**), designed to assess the reliability of per-sample base calls and minimise false positives, including multi-allelic sites and private SNPs likely arising from environmental contamination. We additionally excluded three previously published ancient genomes based on the presence of singleton or clustered mutations likely deriving from non-*Yersinia* species (**Supplementary Note 3, Supplementary Figures 1-4**). This pipeline yielded a set of 2,804 high-confidence SNPs across 288 *Y. pestis* genomes (84 ancient DNA and 204 modern DNA), which were used for maximum likelihood phylogenetic reconstruction (**Supplementary Tables S6-7, Supplementary Figure 5-7**). To contextualise genomic placement within the broader *Y. pestis* phylogeny, we conducted a time tree analysis (**Supplementary Figures 8-10, Supplementary Note 4**) and additionally assigned genomes to likely historical outbreaks aided by a previous study which applied phylogenetically informed radiocarbon modelling (PhIRM) to Second Plague Pandemic *Y. pestis* genomes^20^ (**Supplementary Note 5, Supplementary Table S8**).

All 14 *Y. pestis* genomes that were mapped at >5x mean coverage fell phylogenetically within *Y. pestis* Branch 1 (**Figure 2**), spanning multiple sub-lineages associated with the Black Death and subsequent Second Plague Pandemic outbreaks. Four genomes clustered directly within the Black Death lineage. These included three identical genomes from Lund in Sweden: LUN252 (64.6x), LUN124 (23.0x), and LUN105 (62.1x), and a fourth observation from Lund which differed from the others by only a single SNP (LUN106, 52.4x).

**Figure 2:**
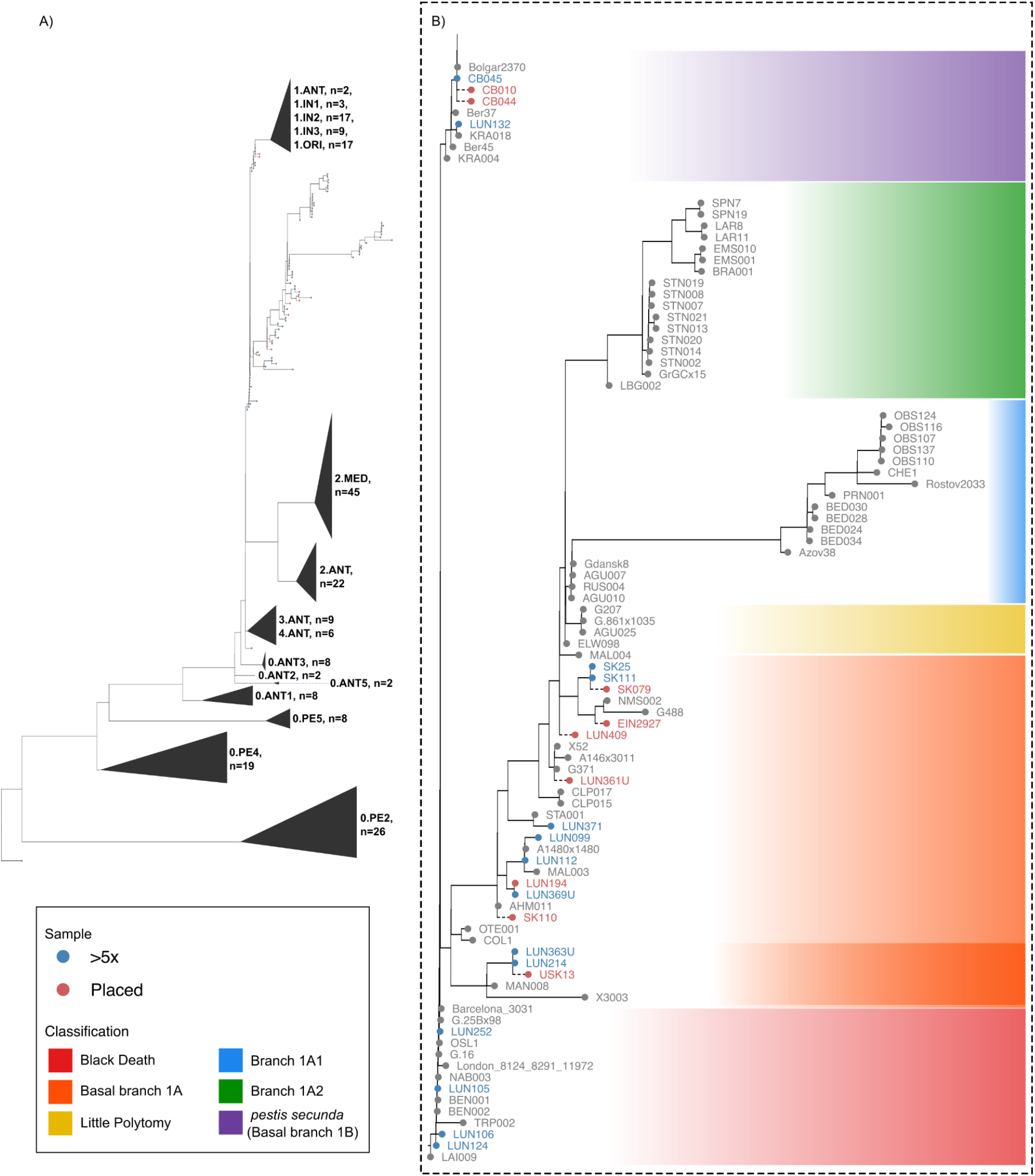
Maximum likelihood phylogeny across the *Y. pestis* dataset. A) Dataset wide maximum likelihood phylogenetic tree. B) Expanded subset of the phylogeny depicted in A highlighting phylogenetic clades involved in the Second Plague Pandemic. *Y. pestis* genomes from this study, and publicly available data that had high enough coverage to be used for constructing the phylogenetic tree, are shown in grey and blue respectively. Lower coverage samples that were placed into the phylogeny are shown in red. The branch length of placed genomes (dashed lines) indicates topological position rather than branch length. Coloured background shading indicates which pandemic phase, and previously defined phylogenetic clade, samples are assigned to. The darker orange shading within basal branch 1A highlights the sub-branch discussed in text.

Eight of the 14 genomes recovered at >5x fell within the diversity of basal branch 1A. This included LUN369U (40.0x) and LUN099 (26.7x), as well as LUN112 (6.6x), which were closely related to *Y. pestis* genomes MAL003 (Estonia), A1480×1480 (Denmark) and AHM011 (the Netherlands)^20,55^. Previous analysis of the human DNA of LUN099 and LUN369U suggests that these individuals were full siblings estimated to be 15-20 years and 7-8 years in age at time of death respectively^44^. The *Y. pestis* infections they carried however differed by at least five SNPs, suggesting these infections derive from independent outbreaks over these possible periods, with LUN369U likely associated with the described outbreaks in Sweden dating to c.1404-1405, 1413, 1421-1422 or 1430-1431^35,56–58^ and LUN099, most likely an infection during the outbreak of 1451-1452^35^ (see **Supplementary Note 5**). LUN112, identical to a genome from Horsens, Denmark (A1480×1480), is a likely representative of outbreaks in the early to mid 15th century (c.1413, 1421-1422 or 1430-1431). We additionally recovered two identical *Y. pestis* genomes in Trondheim from individuals SK25 (46.6x) and SK111 (42.8x), estimated to share a common ancestor in approximately 1521-1540, which are basal to a clade including the previously published genomes NMS002 and G488 from Cambridge (United Kingdom) and Latvia^21,59^ and which are likely to be associated with one of several documented epidemics in the region in the late 15th century.

The remaining two reconstructed high-coverage genomes fell within the diversity of the *pestis secunda* branch 1B. LUN132 (73.6x) is identical to the published genome KRA018 (Germany)^18^ and likely derived from the 1359 *pestis secunda* outbreak in Lund^36,42,43^. CB045 from Armenia (5.4x) is identical to the published genome Bolgar 2370 (Russia)^22^, falling in the *pestis secunda* clade. These additional *Y. pestis* genomes from the *pestis secunda* expand the geographic range of genetically characterised infections. Furthermore, our genome from Armenia reinforces a likely eastward trajectory of the *pestis secunda*, filling in a geographic gap between Central European samples (KRA018, KRA004) and the Bolgar2370 genome recovered from Tatarstan^18,22^ (in the present-day Russian Federation) linked to the Volga region outbreak of 1364^60^.

An interesting feature of our dataset is the concentration of *Y. pestis* genomes recovered from Lund over a period of approximately 270 years, in contrast to many ancient *Y. pestis* studies that consider genomes pooled across multiple different sites. This provides an opportunity for longitudinal assessment of the epidemic characteristics of *Y. pestis* present at a single location. Over this period, we identify *Y. pestis* infections in Lund which could be placed in seven distinct phylogenetic clades across branches 1A and 1B, with a median pairwise phylogenetic distance of ∼15.7 substitutions among high-coverage genomes, compared to ∼18.35 across all Black Death and basal branch 1A genomes. The phylogenetic diversity observed in Lund is not significantly different from the levels of diversity observed across all currently available high coverage Black Death and basal branch 1A genomes considered in our analysis (permutation test; n = 100000; Δmedian = −2.65; p = 0.136). These findings support a model in which plague repeatedly recurred in the region through successive epidemic waves introduced from elsewhere, rather than being maintained primarily through continuous transmission from a persistent local reservoir.

### Phylogenetic placement of lower coverage *Y. pestis* genomes from the post-Black Death period

Ancient DNA datasets frequently include genomes of variable coverage. In this study, we recovered data from an additional 18 individuals with some evidence of *Y. pestis* infection that had genome-wide coverage below our inclusion criteria for conventional phylogenetic analysis (**Supplementary Tables S1-3**). In order to leverage those *Y. pestis* genomes recovered at between 1-5x mean coverage, we applied a workflow built upon the phylogenetic placement algorithm pathPhynder^61^ (see **Methods, Supplementary Note 6, Supplementary Tables S9-11**). In short, pathPhynder allows low coverage genomes to be placed into a reference phylogeny through evaluation of allelic support at phylogenetically informative sites, accounting for missing data and post-mortem ancient DNA damage. Downsampling analysis indicated that this approach could reliably place *Y. pestis* genomes with coverage as low as ≈0.7x, although placement accuracy depended on the local tree structure (**Supplementary Note 6, Supplementary Figure 11**).

Using this approach, we confidently assigned nine additional ancient genomes to established *Y. pestis* lineages, including three from Lund, two from Trondheim, two from Armenia, one from Eindhoven, and one from Iceland (**Figure 2**). We found that our lower coverage genomes from Lund (LUN194; 0.7x) and Trondheim (SK110; 2.2x) fell with previously published genomes across central to northern Europe (AHM011 Netherlands, MAL003 Estonia and A1480×1480 Denmark) and high-coverage genomes LUN369U, LUN099 and LUN112. Similarly, LUN409 (3.8x) was placed with the Estonian strain MAL004^20^. We found that SK079 (Trondheim, 1.9x) and EIN2927 (Eindhoven, 2.5x) were assigned to the same Branch 1A clade including additional high-coverage genomes from Trondheim (SK25, SK111) and *Y. pestis* genomes from Cambridge (NMS002)^21^ and Latvia (G488)^59^.

Interestingly, we found that the Icelandic USK13 genome could be confidently placed within a previously under-characterised sub-lineage of basal Branch 1A, representing the first direct genomic evidence of *Y. pestis* in Iceland (**Figure 2**). This clade also includes two *Y. pestis* genomes we recovered from Lund (LUN363U, LUN214), MAN008 recovered from a mass grave at Manching-Pichl in Germany^21^ and X3003 from Sejet (Horsens) in Denmark^62^. Through molecular dating, we estimate that the most recent common ancestor of this clade dates to 1375-1385 (**Supplementary Figure 10**). The Danish genome X3003, recovered from an individual radiocarbon-dated to 1490-1646, suggests that this clade of *Y. pestis* persisted for at least a century following its estimated diversification in the late fourteenth century. Together, its identification in Germany, Denmark, Sweden and Iceland indicates that this clade was both geographically widespread across northern Europe and the North Atlantic and maintained over an extended period during the Second Plague Pandemic.

Historical records document only two plague outbreaks in Iceland, in 1402-1404 and 1494-1495, with the latter thought to be imported from England^63^. Radiocarbon dating of USK13 (calibrated date: 1434-1513 calAD (95.4%) and recalibrated date: 1460-1814 calAD (95.4%)) after taking into account the marine reservoir effect is more consistent with the later outbreak. Relative to the radiocarbon-dated Lund genomes from this lineage (LUN363U and LUN214), USK13 carries nine additional nucleotide substitutions, seven confidently supported at ≥3x coverage, including two predicted non-synonymous substitutions in the genes *rcsC*^64^ (Ser5Phe) and *lptG*^65,66^ (Gln137Arg). Assuming a mutation rate of approximately one substitution every 4.8 years^20^, this suggests an estimated date of USK13 of 1436-1503, again more consistent with the later outbreak, noting that this date is, if anything, likely to be an underestimate since we are unable to account for missing SNPs located on the terminal branch leading to USK13 due to low coverage.

Finally, we found that two low coverage genomes from Armenia, CB010 (1.41x) and CB044 (1.50x) could be robustly placed as additional *pestis secunda* strains together with the high coverage Armenian CB045 and Bolgar 2370 from Tatarstan (**Figure 2**). The placement of these genomes is consistent with these infections deriving from the 1364-1365 outbreak documented in Armenia^67^, further reinforcing the eastward dissemination of the *pestis secunda* before its apparent disappearance from Europe.

### Identification of mutational signatures defining the Black Death and the *pestis secunda Y. pestis* waves

To characterise the accumulation of chromosome-borne mutations across the Black Death and subsequent outbreaks, we applied a framework we term Ancestral Reference Mapping (ARM), which reconstructs mutations along the phylogeny relative to their most recent common ancestor.

We first applied ancestral state reconstruction to infer the most parsimonious ancestral genome at the node giving rise to branches 1-4, otherwise known as the Great Polytomy (see **Methods, Supplementary Figure 12**). This reconstructed Great Polytomy genome was identical, at covered sites, to the previously published ancient genome BSK001^19^ from Kara-Djigach which falls immediately basal to Black Death strains, providing independent validation of the inferred ancestral sequence. We then used this reconstructed genome as the reference for read mapping and variant calling, rather than CO92, which as a modern strain, represents a derived reference already carrying mutations that would not have been present in our ancient genomes. Following variant calling, we performed ancestral state reconstruction to assign a permissive set of substitutions and short insertions and deletions (indels) to specific phylogenetic branches of the *Y. pestis* phylogeny (see **Methods, Supplementary Table S12-14, Supplementary Figure 12**). We applied a relaxed filtering criteria to these SNPs and indels to retain homoplastic mutations, as these may reflect convergent evolution and are therefore of potential evolutionary interest (see **Methods**).

The ARM approach provided a temporally resolved view of the mutational changes across post-Black Death lineages since their most recent common ancestor. In total, we identified 16 unique mutations at the nodes leading from the Great Polytomy to the most basal node of the *pestis secunda*, comprising five substitutions and 11 indels in both genic and intergenic regions (**Supplementary Table S13**). We noted one intergenic mutation upstream of YPO_RS09215 (1871476, A->G) which showed a phylogenetically informative pattern of being acquired and maintained in descending lineages, while others were more homoplastic across the tree (**Supplementary Figure 13**). Upon excluding mutations in intergenic regions, six non-synonymous mutations remained (**Supplementary Figure 14**). We noted that of these six, three exhibited a clear pattern throughout the phylogenetic tree, in which they were acquired and subsequently found in descending lineages, both early in the Black Death and in the *pestis secunda* (see **Figure 3**). The remaining three mutations were homoplastic across branch 1A and the phylogeny. These include a +1T insertion (Ile457fs) in YPO_RS1940, encoding a neuraminidase-like domain protein; a +1T insertion (Asn104fs) in YPO_RS17990, associated with metal ion binding and carbohydrate metabolism; and a -1A deletion (Phe4fs) in YPO_RS10230, a putative DNA-binding protein.

**Figure 3:**
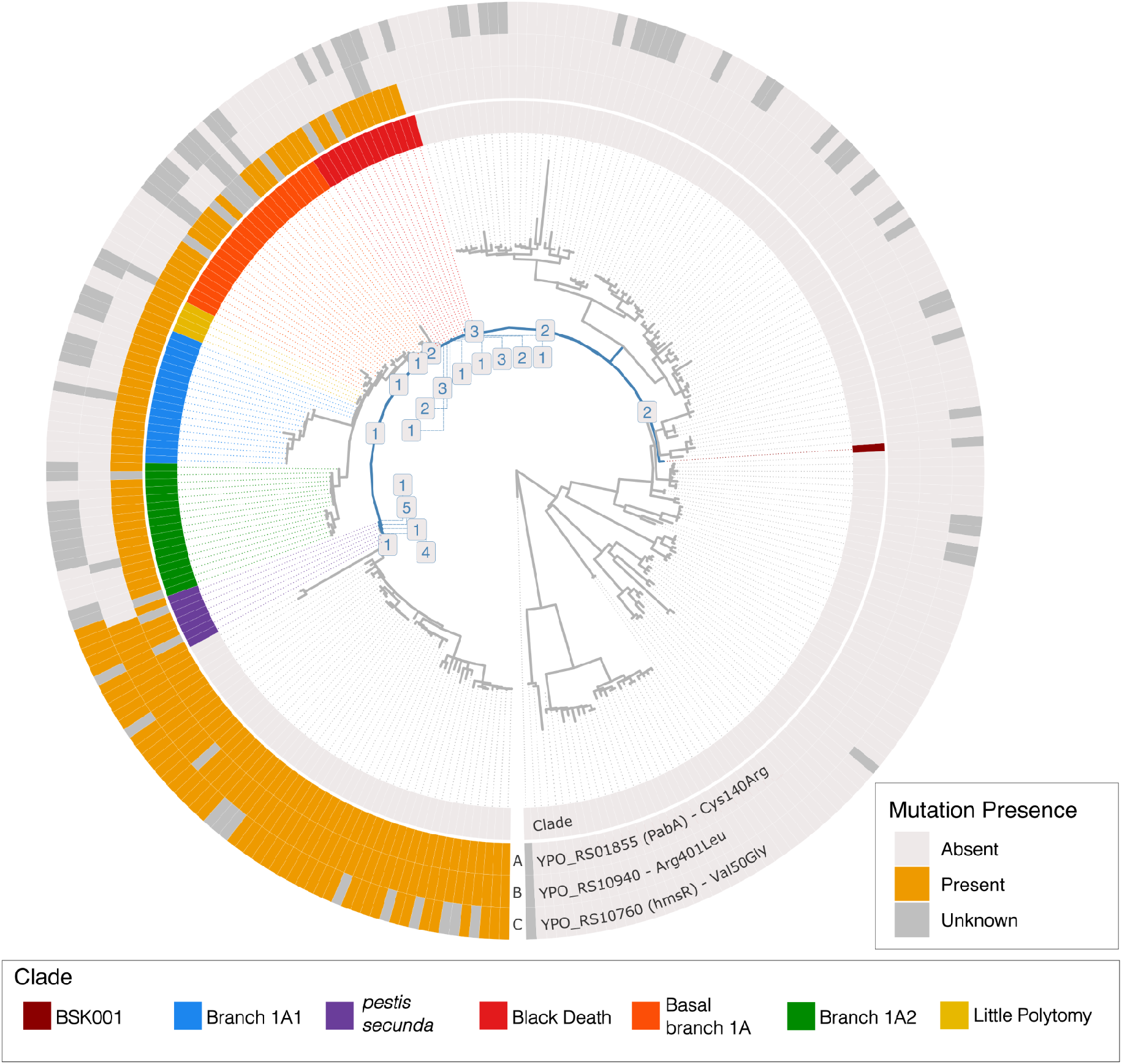
Radial phylogeny and heatmap depicting the presence and absence of three non-synonymous mutations relative to an inferred ancestral reference. Radial maximum likelihood *Y. pestis* phylogeny. The path from the Great Polytomy node to the most basal *pestis secunda* node is shown in blue with the number of mutations assigned to each node labelled. Columns of the heatmap correspond to: the innermost ring is a coloured heatmap indicating the phylogenetic clade as per the legend at bottom; A) the T->C mutation in the YPO_RS01855 (*PabA*) gene; B) the G->T mutation in the YPO_RS10940 gene and C) the T->G mutation in the *hmsR* gene. Colours denote the presence and absence of mutations, with “unknown” characterising sites without sufficient coverage.

We next characterised the potential impact of the three non-synonymous mutations that were acquired and maintained at key transition points in the epidemiology of early Second Plague Pandemic *Y. pestis* (**Figure 3**). None of these mutations has been observed in data screened from the Late Neolithic Early Bronze Age^68^, First Plague Pandemic or extant, basal *Y. pestis* clades^4,5,69,70^ (see **Methods**). Focusing on the early Black Death period we identify a T-to-C transition in YPO_RS01855, resulting in a Cys140Arg change in the protein (**Figure 3**, A). This mutation is found in all post-Black Death lineages (including the globally distributed modern Branch 1B) represented in our dataset with sufficient coverage at this position. This mutation has previously been characterised as a cytosine-to-thymine mutation in BSK001 relative to the CO92 reference genome which carries the derived allele^19^. YPO_RS01855 encodes aminodeoxychorismate synthase component II (PabA), which functions in complex with PabB to produce 4-amino-4-deoxychorismate (ADC), a key intermediate in the biosynthesis of p-aminobenzoate (PABA) and folate. This pathway is essential for bacterial growth and is targeted by sulphonamide antibiotics^71^. Residue 140 is located on a solvent-exposed surface of the PabA protein, away from the active site (residues Cys79, His172, Glu174)^72^ according to the AlphaFold3 model (AF3; pTM = 0.96; **Figure 4**, A-B). Prediction of protein-protein interaction interfaces and modelling of the PabA-PabB complex (pTM = 0.93, ipTM = 0.94) suggest that this residue is unlikely to be directly involved in protein interactions. However, stability ΔΔG predictions suggest that position 140 is located within an energetically constrained region, where the Cys-to-Arg substitution may induce local conformational changes associated with the introduction of a positively charged residue at a solvent-exposed site (**Figure 4**, A-C; **Supplementary Table S15**)^73,74^.

**Figure 4:**
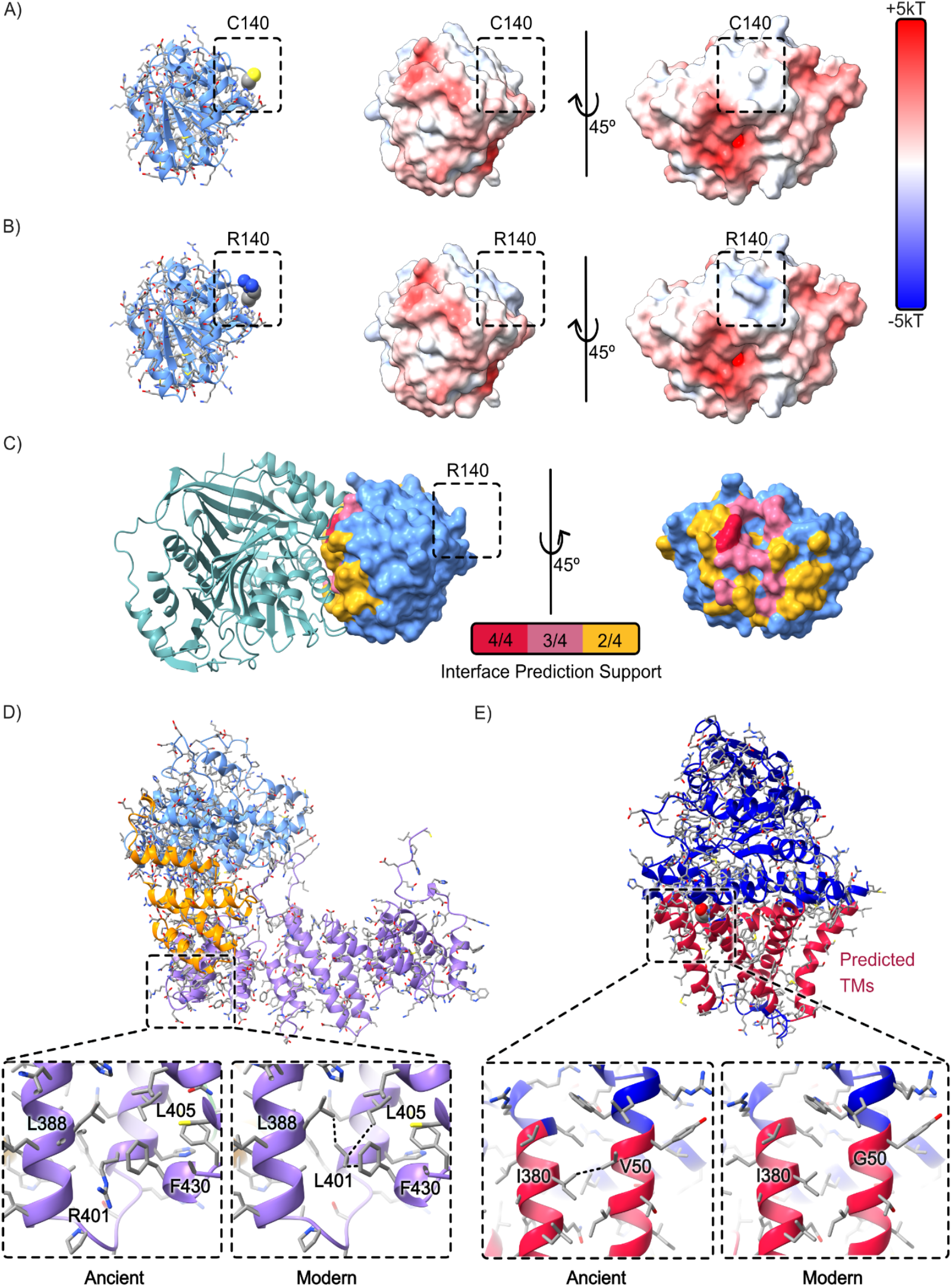
Protein modelling and structure-based analyses localise missense variants that emerged during the Black Death and the *pestis secunda* and are present in present-day *Y. pestis* Branch 1 strains. A) Alphafold3 model of *Y. pestis* ancient PabA (YPO_RS01855; Cys at position 140). B) Alphafold3 model of CO92 PabA (YPO_RS01855; Arg at position 140). A-B) Middle-right panels: surface rendering of the modelled proteins coloured by electrostatic potential (Scale bar: red, -5kT; blue, +5kT). C) Left panel: Interaction complex between PabA (YPO_RS01855; blue surface representation) and PabB (YPO1773; turquoise cartoon representation); Right panel: Surface representation of PabA. Predicted interfacial residues of PabA are coloured based on the number of supporting algorithms: red (4/4), violet (3/4) and yellow (2/4). D) Alphafold3 model of *Y. pestis* YPO_RS10940 (protein domains 1-287, 288-378, 379-703 are coloured in blue, yellow and purple respectively), closed-up views show residue 401 and the hydrophobic contacts mediated by Leu401 in the modern version of YPO_RS10940 (CO92). E) Alphafold3 model of *Y. pestis* hmsR, predicted transmembrane regions are coloured in red, closed-up views show residue 50 and the hydrophobic contacts mediated by Val50 in the ancient version of hmsR.

Two additional non-synonymous mutations arose along the lineage leading to the *pestis secunda* and modern Branch 1B. The first is a G-to-T transition in YPO_RS10940 (Arg401Leu) which was acquired during the *pestis secunda* and is present in all the high coverage *pestis secunda* genomes to date aside from KRA001. It has previously been identified as a defining variant of this lineage^18^ (**Figure 3**, B). YPO_RS10940 encodes a membrane-associated protein implicated in protein binding, although its precise function remains unclear. Modelling of the YPO_RS10940 protein structure with Alphafold3 (pTM: 0.81) shows that position 401 is located within an ɑ-helical region stabilized by hydrophobic contacts. The Arg-to-Leu substitution introduces a hydrophobic residue that is predicted to establish additional intramolecular hydrophobic contacts with Leu388, Ile405, and Phe430, consistent with the predicted stabilizing effect of the substitution (**Figure 4**, D; **Supplementary Table S15**).

The second mutation is a T-to-G substitution in hmsR (Val50Gly). *hmsR* encodes an inner membrane protein within the hmsHFRS operon, which plays a key role in biofilm formation and flea-borne transmission in *Y. pestis*^75–78^, as well as modulating virulence in other gram-negative bacteria^79,80^. This mutation is present in all the *pestis secunda* genomes in our dataset with sufficient coverage, and is present in descendant lineages, including those associated with the Third Plague Pandemic in our dataset (see **Figure 3**). This mutation has not previously been discussed because the modern reference strain CO92 carries the derived allele relative to our reconstructed Great Polytomy (ancestral) reference. In addition, *hmsR* is frequently masked in genomic analyses of *Y. pestis* owing to its previous classification as a non-core region^81^, which may have contributed to this variant being overlooked. However, *hmsR* is consistently represented in our dataset (**Figure 3**). Furthermore, a BLAST analysis of 1272 published Branch 1 *Y. pestis* genomes with complete or near-complete assemblies (≥95% of the CO92 reference length; NCBI, accessed June 2026) found that *hmsR* is present in approximately 92% of genomes, suggesting that its designation as a non-core locus may warrant reevaluation (**Supplementary Table S16**).

Given its important role in biofilm formation in *Y. pestis*, site-directed mutagenesis experiments have been conducted on *hmsR* though the upstream region has not been well characterised^77^. Modelling of the hmsR protein structure with AlphaFold3 (pTM: 0.93), together with transmembrane helix prediction, indicates that the *pestis secunda* mutation Val50Gly is located near the end of the second predicted transmembrane helix. In the model, Val50 is exposed to the lipid bilayer and establishes a hydrophobic interaction with Ile380 in a neighbouring transmembrane helix (**Figure 4**, E). These observations suggest that the Val50Gly substitution could alter local protein stability and dynamics by disrupting hydrophobic interactions with the lipid environment and adjacent transmembrane regions. These observations raise the possibility that this mutation, acquired at the *pestis secunda*, may influence biofilm formation or related regulatory dynamics in *Y. pestis*, although experimental validation will be required to assess functional significance.

## Discussion

*Y. pestis* is one of the most historically consequential zoonotic pathogens, responsible for multiple pandemics that reshaped human populations across continents. Here, we present newly reconstructed *Y. pestis* genomes spanning the Black Death, the *pestis secunda*, and subsequent outbreaks of the Second Plague Pandemic from Sweden, Norway, the Netherlands and Armenia. We also provide the first direct molecular evidence of plague in medieval Iceland. By combining longitudinal sampling from the major cultural and trade centres of Trondheim and Lund with this broader geographic coverage over Europe and the Caucasus, we map the historical phylogeography of *Y. pestis* in human populations, and provide support for an eastward dissemination of the *pestis secunda* consistent with the historically documented epidemic wave.

Despite rich historical documentation and an expanding palaeogenomic record, key aspects of plague persistence, geographic spread, and diversification following the Black Death remain only partially resolved. Integrating genomic data with historical sources allows direct comparison between phylogenetic patterns and recorded outbreaks^20^. The genomes from the time of the Black Death are consistent with a model of rapid clonal expansion: multiple genomes from Lund are identical to other genomes from the Black Death (LUN105, LUN252, LUN124), reflecting the low diversity characteristic of the initial pandemic wave.

In contrast, the post-Black Death period is marked by increased phylogenetic complexity. The *pestis secunda*, the first major epidemic wave following the Black Death, has been proposed to have spread from Germany in a concentric circle-like expansion before disappearing from Europe, with descendant lineages dispersing into east Africa and eastward towards China, where they later gave rise to extant Third Pandemic lineages^28,82^. Despite growing interest in the *pestis secunda*^18,28,60,83^, genomic observations remain limited. Our analyses provide support for this model. We identify four new *pestis secunda* genomes, including LUN132 from Lund, the most northerly genomic observation to date, which is identical to a German genome (KRA018)^18^. In Armenia, we recovered three *Y. pestis* genomes likely associated with the 1364-1365 *pestis secunda* outbreak^60^, including a high-coverage genome identical to Bolgar2370 from the Middle Volga region^22^.

Together, these new *pestis secunda* genomes reinforce a rapid spread of the epidemic from South-Central Germany from 1356 on^18,28^, reaching as far east as Armenia and Bolgar in less than 10 years. Additionally, these data suggest different sub-branches (reflecting different geographic routes of spread) that the *pestis secunda* was split into: the first (terminal) leading to KRA018 and LUN132, and the second forming another branch containing Bolgar2370 and CB045. This lineage subsequently disappeared from Europe but its descendants likely gave rise to 1.ANT branches thriving in East Africa^84^, 1.IN branches circulating in western China and Tibet and 1.ORI - the causative lineage of the Third Plague Pandemic, which presumably emerged in the Yunnan province of China in 1772, and spread globally from 1894 onwards^12^. The geographic locations, and potential hosts, in which *Y. pestis* persisted in the ∼400 years between the end of the *pestis secunda* and the emergence of the Third Plague Pandemic still remain unknown, although multiple reservoirs and migration routes have been suggested^82^.

Our work provides the first genomic evidence of *Y. pestis* in Iceland, addressing longstanding uncertainty over whether *Y. pestis* was the causative agent of the historically documented Icelandic Second Plague Pandemic outbreaks, in part due to the apparent absence of rats from Iceland during this period^38,39^. USK13 differs by nine SNPs from the Swedish genomes LUN214 and LUN363U and falls within a previously under-characterised sub-lineage of basal branch 1A. This lineage appears to have been long-lived and spatially extensive, with further representatives from Germany (MAN008; 1283-1390) and Denmark (X3003; 1490-1646). Relative to the Lund genomes, the Icelandic USK13 carries two non-synonymous substitutions: a Ser5Phe substitution in *rcsC*, which encodes the membrane-bound protein involved in the Rcs phosphorelay pathway which helps regulate biofilm formation^64^, and Gln137Arg in *lptG*, which encodes the lipopolysaccharide export ABC transporter permease LptG involved in maintenance of the cell wall^65,66^. Although the functional consequences of these substitutions remain unknown, it is unlikely that they represent local diversification or adaptation within Iceland, where plague is only documented in two short outbreaks in 1402-1404 and 1494-1495. Nevertheless, whether this lineage acquired specific adaptations prior to its arrival in Iceland is of particular interest given the unusual ecology of plague in this setting and the possibility of transmission mechanisms that did not depend on rats or other rodent hosts^85–87^. The genomic identification of *Y. pestis* in Iceland therefore provides an important new perspective on this debate, shifting the question from the identity of the causative agent towards how *Y. pestis* was transmitted and maintained in Iceland. Further genomic observations of this currently sparsely sampled lineage may help resolve its geographic history and provide additional insight into the reservoir ecology and transmission dynamics of plague in Iceland.

Given the clonal nature of *Y. pestis*, SNP-based characterisation provides a powerful framework for phylogenetic placement. Unlike more rapidly evolving pathogens, the clonality and slow substitution rate of *Y. pestis*, combined with the rarity of back mutations, means that individual SNPs can serve as reliable phylogenetic markers, enabling consistent clade assignment across datasets. This is particularly valuable in ancient DNA studies, where genome coverage is often incomplete. Here, we apply a reproducible mutation-filtering pipeline to identify phylogenetically informative SNPs that recover the established *Y. pestis* phylogeny across the Second Plague Pandemic.

Building on this, we developed a mutation-mapping framework that departs from conventional variant annotation against the CO92 reference genome, which was isolated in 1992 and hence represents a derived Third Pandemic lineage^88^. Instead, we reconstructed the probable ancestral sequence at the Great Polytomy and used this as the reference for read mapping and variant calling. This allowed us to catalogue non-synonymous changes arising along major Second Plague Pandemic lineages relative to their ancestral state, providing a more biologically meaningful account of evolutionary change during the early pandemic. Notably, we identified three non-synonymous mutations acquired between the Great Polytomy and the *pestis secunda*, all of which are seen in Third Plague Pandemic lineages. Two of these mutations have been previously under-characterised because CO92 carries the derived allele and/or the affected locus has been masked from previous phylogenetic alignments^81^. These observations highlight how ancestral reference mapping can recover both the occurrence and direction of evolutionary transitions that may be overlooked or misinterpreted when ancient genomes are analysed solely against a derived modern reference.

Two of the non-synonymous mutations identified fall in genes with known and potentially relevant functions in *Y. pestis* biology. The first, acquired in the Black Death, affects *pabA*, which is involved in folate biosynthesis, a pathway essential for nucleotide production and bacterial replication, and an established antimicrobial drug target^71^. This gene, and it’s counterpart *pabB*, have also been implicated in the fitness of *Y. pestis* in deep-tissue infections^89^. The second, acquired in the *pestis secunda*, affects *hmsR*, which encodes an inner-membrane protein involved in extracellular biofilm matrix production. This is of particular interest given the critical role biofilm formation plays in *Y. pestis* transmission, where blockage of the flea proventriculus increases biting frequency and enhances transmission efficiency through regurgitation of infected blood^75–78^. Structural modelling predicts that both mutations, while outside of the active site of their respective proteins, may still alter protein confirmation and stability. As such, it is unclear whether these substitutions contributed to the epidemiological success of their respective lineages or simply coincided with major transitions in the presentation of *Y. pestis*. Nevertheless, the emergence of non-synonymous changes in genes with plausible biological relevance at the Black Death and the *pestis secunda* suggests that their functional consequences warrant further investigation.

It is interesting to speculate on whether adaptive changes in *Y. pestis* were necessary for the epidemiological shifts in Branch 1 prior to the Black Death and the *pestis secunda*, or whether changes in host populations and pathogen ecology were sufficient to drive these transitions. With the (re-)introduction of plague into Europe during the Black Death, *Y. pestis* may have been able to establish new reservoirs, outside of the birthplace of the causative lineage in Central Asia. By the late 14th century, European populations had experienced substantial exposure to plague, potentially altering population-level susceptibility to subsequent outbreaks. Prior infection may have conferred some degree of immunity; for example, today antibodies to the major F1 antigen have been shown to persist for more than a decade following infection^90^, though the extent to which this corresponds to long-term protective immunity is uncertain. Demographic turnover would, however, have continuously replenished the pool of immunologically naïve individuals, particularly among those born in the intervening years; a cohort that may have been disproportionately susceptible, potentially contributing to the elevated mortality among younger individuals recorded by chroniclers during the *pestis secunda*^29^. However, comparisons with mortality during the Black Death, when populations were also largely unexposed, suggest that variations in prior exposure alone are unlikely to explain the apparent differences in the lethality, spread, and demographic impact between successive outbreaks^91^. The interplay between pathogen evolution, host immunity, and reservoir ecology thus remains an important and unresolved area for future work.

Our work provides new insights into the evolutionary and epidemiological dynamics of *Y. pestis* during and after the Black Death and the *pestis secunda*. By integrating genomic data with historically documented outbreaks, we refine existing evidence for repeated reintroduction of plague into northern Europe and the Caucasus and extend the phylogenetic diversity present within Black Death and post-Black Death epidemic waves. We established a previously poorly characterised early emerging sub-lineage within basal branch 1A that persisted for at least a century and was geographically widespread across northern Europe and the North Atlantic. Its identification in Iceland provides the first direct genomic evidence of *Y. pestis* in the country and implicates this lineage in one of the two historically documented Iceland plague epidemics, providing new context for questions surrounding plague transmission in the apparent absence of rats. In addition, our *Y. pestis* genomes from Armenia strengthen the case for a rapid eastward spread of the *pestis secunda* across Eurasia prior to its disappearance from Europe. Through an ancestral reference mapping approach, we document non-synonymous mutations defining the evolutionary trajectories of the Second Plague Pandemic and in doing so note structural changes in genes implicated in metabolism and biofilm formation at the transition points of the Black Death and the *pestis secunda*. Collectively, our findings demonstrate how integrating genomic, historical, and evolutionary perspectives can resolve the movement, persistence and diversification of *Y. pestis* across successive epidemic waves and provide new insight into the changing dynamics of the Second Plague Pandemic.

## Materials and Methods

### Sample Processing, DNA Extraction and Library Preparation

The majority of the Trondheim and Lund DNA data analysed derives from Liu et al^44^, where full methodological information on the data generation can be found. For the remaining samples DNA was extracted following a modified version of Moreno-Mayar *et al*. 2018’s^92^ protocol as described in Liu *et al*.^44^ (**Supplementary Note 2**). Extracts were then converted into single-strand DNA Illumina libraries using the Santa Cruz Reaction (SCR) protocol of Kapp *et al*. 2021^93^. Libraries for most of the samples were initially prepared without UDG treatment to assess DNA damage profiles (**Supplementary Table S2**). These libraries were screened for the presence of pathogen DNA using MALT^52^ (v0.4.1) and the MaltExtract and AMPS tools integrated in the HOPS^53^ pipeline (**Supplementary Table S3**). Based on the pathogen screening results, 32 individuals were chosen for *Y. pestis* whole genome capture (for full details please see **Supplementary Note 2**).

### Dataset Curation

Previously published aDNA *Y. pestis* genomes were identified using the “ancientsinglegenome-hostassociated” samples table from AncientMetagenomeDir^94^ (v23.09.0), accessed 01/10/2023. The table was then imported into Julia^95–97^ (v1.10.0) and filtered for *Y. pestis* aDNA genomes that have been confidently linked to the Second Plague Pandemic^7,17–22,33,55,59,62,98–101^. The resulting table was input to the AMDirT^102^ (v1.4.6) convert command line tool to generate a download script for the raw sequence data (Fastq) and generate the nf-core eager input file^103^.

### Processing Genomic Data

Raw sequencing data were processed using the nf-core eager pipeline^103^ (v2.4.7) implemented in Nextflow^104^ (v22.10.0). Adapters were removed using AdapterRemoval^105^ (v2.3.2) with the minimum adapter overlap set to 3. Additionally, samples that were sequenced with a two-colour chemistry had poly-G tails removed using fastp^106^ (v0.20.1). Ancient libraries were mapped using BWA aln^107^ (v0.7.17) with a range of parameters depending on what level of UDG treatment they had (**Supplementary Note 3**). Where necessary, the resulting library bam files were merged using samtools^108^ (v1.12) merge, deduplication was performed using Picard MarkDuplicates^109^ (v2.26.0) and the bam file was then sorted and indexed using samtools.

Modern genomes were mapped using BWA mem with default parameters where raw sequencing data was available. For other genomes available as contigs or consensus sequences, short reads were simulated as described in **Supplementary Note 3**. Resulting bam files were filtered using samtools, requiring a mapping quality of 37; unmapped reads were discarded, and deduplication was performed using Picard MarkDuplicates. Genotyping was performed using GATK Unified Genotyper^110^ (v3.5.0) with the EMIT_ALL_SITES flag. SNP tables and alignments were generated using MultiVCFAnalyzer^111^ (v0.85.2). A custom pipeline of filters to remove erroneous calls due to damage and environmental contamination were then applied (**Supplementary Note 3**). The final SNP calls are provided in **Supplementary Table S6**.

### Phylogenetic Reconstruction

The filtered SNP alignment was used to construct maximum likelihood trees in IQTree2^112^ (v2.4.0). The ultra-fast bootstrap and SH-aLRT parameters were set to 1000 replications, and the collapse near-zero branches flag was passed. GTR+F+ASC+R2 was used as a substitution model. Visualisations of the trees were made in R^113^ using the ggplot2^114^ (v3.4.4) and ggtree^115^ (v3.10.0) packages. A matrix of SNP distances between genomes was calculated using snp-dists, and the distance between genomes within clades was visualised as heatmaps in Julia using Makie.jl^116^ (v0.20.4) and CairoMakie.jl (v0.11.5) (**Supplementary Figures 5-7, Supplementary Table S7**).

To estimate the temporal behaviour of the phylogeny, a maximum likelihood tree of branch 1A and the *pestis secunda* was generated with IQTree2. The phylogenetic tree and associated tip dates were imported into TempEst (v1.5.3) and the root-to-tip correlation was assessed to confirm a temporal signal^117^. A time calibrated phylogenetic tree was then generated with BactDating (v1.1.2)^118^ using a strict gamma model with a coalescent tree prior and the update root flag set to false. After 5e^+6^ Markov chain Monte Carlo (MCMC) iterations, the effective sample sizes and traces of the parameters were interrogated to ensure MCMC convergence (see **Supplementary Note 4, Supplementary Figures 8-10**).

To estimate the distances between nodes (internal and terminal) in terms of estimated mutation counts when accounting for placed samples (see **Phylogenetic placement**), a set of nine modern genomes was randomly selected from across the phylogenetic tree. Modern genomes were chosen because they typically contain far fewer missing or ambiguous SNP calls compared to aDNA samples. The distances between each combination of nodes on the tree were then calculated using the Phylo.jl^119^ (v0.5.2) package in Julia, and the SNP distances between the pairs were extracted from **Supplementary Table S7**.

For each pair of genomes, a scaling factor was determined by dividing the SNP distance between the genomes by the path length between the nodes. The final conversion factor was obtained by averaging these pair-specific values, which provides a tree-wide estimate that is less sensitive to noise or rate variation affecting any single pair. This average scaling factor was then applied to any branch or node-to-node distance in the tree by multiplying the phylogenetic distance by the scaling factor to obtain an estimated number of mutations.

### Phylogenetic placement

The filtered SNP table from MultiVCFAnalyzer (see **Supplementary Note 3**, **Supplementary Table S9**) was converted to a VCF file and, along with the phylogenetic tree (see **Phylogenetic reconstruction**), was used to identify diagnostic SNPs with the tool Phynder (v1.0) (**Supplementary Table S9**). Placement was then performed with pathPhynder^120^ (v1.2.3), specifying the best path method, providing the output file from Phynder with the filtered and deduplicated bam files that had a mean coverage of less than 5x. Placement was conducted with the minimum base quality set to 30 and the pileup read mismatch threshold set to 0.9 (see **Supplementary Note 6, Supplementary Figure 11**).

### Phylogenetic association with historical outbreaks

To associate different recovered genomes with their most probable historically documented outbreaks, results from the PhIRM (phylogenetically informed radiocarbon modelling) approach as developed by Keller *et al*. 2026 were used^20^. In brief, this method looks to integrate phylogenetic information with Bayesian radiocarbon modelling to juxtapose the modelled date ranges against the documentary record (**Supplementary Table S8**), thus associating the sequenced and analysed genomes with a series (between ∼1-5) of documented plague outbreaks. Based on the contextualisation of genomes presented in Keller *et al*, a full historical contextualisation of new genomes is provided in **Supplementary Note 5, Supplementary Table 8**.

### Ancestral Genotyping Approach

Filtered and deduplicated bam files with a mean coverage greater than 5x were selected. Samtools (v1.22.1) mpileup was first used to generate information on the nucleotides covering each position, with the minimum map quality set to 25, the minimum base quality set to 20, and the read group (RG) tags were ignored. Variants were then called with several filters to help mitigate erroneous calls. The first step was to calculate the z-score for the coverage of all positions across the dataset; any site that had a coverage more than two standard deviations above the alignment’s mean coverage was suspected of being due to short fragments spuriously aligning and was excluded. Following this, any position that did not meet the minimum coverage threshold (3x for ancient, 10x for modern) was excluded.

In order to call a site as homoallelic, an allele frequency of 0.9 was required, with a minimum frequency of 0.1 required to be considered a heteroallelic call. Insertions were called if they passed a minimum depth threshold of 5-fold or a z-score of -0.5, whichever was stricter. For a deletion to be called, it would need to pass the same threshold as the insertion; if this occurs, the deletion calls were combined with the calls at the site where the deletion is located and the site was recalled, incorporating the additional information.

Mutations were considered as potentially deriving from post-mortem damage if they occurred in an ancient genome and exhibited a base change typically attributed to aDNA damage (G - > A, C -> T). Whenever such a base change was called, the proportion of observations on the forward and reverse strands was calculated. If the proportion of forward (C -> T) or reverse (G -> A) calls was less than 0.3, then the call was kept. If the proportion of forward or reverse calls was greater than 0.3 but less than 0.8, a minimum depth threshold of 5-fold or a z-score of -1, whichever was stricter, was applied in order to call the site. If the proportion of forward or reverse calls was greater than 0.8, then the call was excluded.

### Ancestral Reference Mapping (ARM) and Mutation Pathway Analysis

To reconstruct a reference genome for the Great Polytomy to apply ancestral reference mapping (ARM), we utilised SNP data (see **Supplementary Note 3**) to generate pseudo-fasta files. These fasta files, in conjunction with our maximum likelihood phylogeny (see **Phylogenetic Reconstruction**), were used to conduct ancestral sequence reconstruction (ASR) using Iqtree2, employing the GTR+F+R2 model (see schematic of workflow in **Supplementary Figure 12)**^112^.

To map the mutations identified through the ancestral variant calling approach back to the tree, we began by converting the output of the ancestral genotyping approach (see section **Ancestral Genotyping Approach**) to a mutation table, using data.table (v1.180.0), matching rows to the tips of the phylogenetic tree. To help reduce the complexity of the character state space, all indels were collapsed down to either “+” for insertions or “-“ for deletions. The resulting mutation table was imported into R along with the phylogenetic tree, assigning names to the internal nodes^121–123^. To reduce the memory requirements of the approach, the mutation table was then split into chunks, and most parsimonious reconstruction ASR was conducted, implemented in phangorn^124^ (v2.12.1). The output was then combined into a single output matrix and mutations were assigned back to the nodes. The path from the Great Polytomy node to the most distal *pestis secunda* node was then calculated. To do so, all mutations assigned to those nodes that were present in 25% or more of the Black Death and the *pestis secunda* samples were extracted. Indels were then re-expanded to their original notations, and the names of the genes in which the mutations occurred were extracted. We additionally assessed whether the identified mutations were fixed in Late Neolithic Early Bronze age (LNBA) data^68^ and Justinianic samples^4,5,69,70^ that were not included in the tree. *In-silico* characterisation of all mutations identified was conducted using SnpEff^125^ (v5.3a) and a custom database constructed using the reconstructed Great Polytomy reference, and gtf/gff annotations were lifted over from the CO92 reference (NC_003143.1) using Liftoff^126^ (v1.6.3) (**Supplementary Tables S11-13**).

### Protein modelling and interface prediction

The protein structure of monomeric PabA (YPO_RS01855), hmsR and YPO_RS10940, and the PabA-PabB interaction complex (YPO_RS01855-YPO1773) was modelled with AlphaFold3^127^. Modelling was performed using the amino acid sequences from the *Y. pestis* CO92 reference strain. Four complementary structure-based tools were used to predict interfacial residues in the YPO_RS01855 AlphaFold3 model: Meta-PPISP^128^, PeSTo^129^, ISPRED4^130^, SPPIDER^131^. Predicted interfacial residues were then ranked based on the number of supporting algorithms (range 0-4).

### *In-silico* prediction of stability changes (ddG) upon mutation

We used Rosetta to estimate how R140C mutation influences the stability of YPO_RS01855 using the AlphaFold3 modelled protein structure^73,74,132^. We used the cartddg2020 protocol to estimate stability ddG^73^. We ran 10 separate rounds of energy minimization (ScoreFunction= fullatom, weights= ref2015_cart, symmetric=0, FastRelax name=fastrelax, scorefxn=fullatom, cartesian=1 repeats=1, mover=fullatom)^133–135^, followed by ddG calculation (iterations: 5, score_cutoff: 1, bbnbrs: 1, weights: ref2025_cart, frag_nbrs 2, ignore_zero_occupancy: false, flex_bb: false, fa_max_dis: 9, legacy: false). Finally, we used a scaling factor of 2.94 to convert Rosetta Energy Units (REU) to kcal/mol^74^.

### Electrostatics

Electrostatics potentials were computed with the finite difference Poisson-Boltzmann method^136^. The minimized Alphafold3 model for YPO_RS01855 (using Rosetta, see above) was used as the initial structure. The R140C mutation was modelled using ChimeraX^137^ using the Dunbrack rotamer library^138^. Atomic models were converted to PQR files using the PDB2PQR server using default options (protonation states at pH 7 were defined by PROPKA, forcefield: PARSE)^139–141^. Generated PQR files were processed with the pyDelphi server (default options; grid scale 2.2 grid points/Å)^142^. Protein surfaces were coloured by the electrostatic potential using ChimeraX.

## Acknowledgements

CF is funded by a LIDo DTP studentship. LvD is supported by a UKRI Future Leaders Fellowship MR/X034828/1. MTPG acknowledges Danish National Research Foundation award DNRF143 and Carlsberg Foundation award CF18-1109 for funding the data generation. LTL is supported by the Danish National Research Foundation through the PandemiX Center (grant no. DNRF170). L.Y. and Z.K. were supported by the Higher Education and Science Committee of the Ministry of Education and Science of Armenia (research project no. 21AG-1F025). PS is supported by an ERC Synergy (#101118880) and UKRI EPSRC Horizon Europe UK Guarantee Funding (EP/Z003288/1) grant awarded for the *Synergy-Plague. Reconstructing the environmental, biological, and societal drivers of plague outbreaks in Eurasia between 1300 and 1900 CE*.

The authors wish to thank the NTNU University Museum, the Norwegian National Committee for Research Ethics on Human Remains, Kulturen Museum Lund (Sweden), and the National Academy of Sciences (Armenia) for permitting and supporting the sampling of the skeletal remains. We thank the Municipality of Eindhoven for providing a permit for the use of human sample material from the St. Catherines churchyard excavation in Eindhoven, the Netherlands. We thank the National Museum of Iceland, and in particular Dr Joe W. Walser III, Curator of Physical Anthropology, who granted access to the Icelandic remains. Ethical approval to generate palaeogenomic data from the ancient Norwegian samples was provided by the Norwegian National Committee for Research Ethics on Human Remains.

We additionally wish to thank Finley Grover Thomas and Ida Moltke for additional analytical support and Jennifer Klunk and Daciel Arbor Biosciences for their expertise developing the capture enrichment approach.

## Code Availability

Custom scripts used for the analyses in this study have been deposited in a public GitHub repository and are available at: https://github.com/theHatIsBack/The-Genomic-Landscape-of-Post-Black-Death-Epidemics-in-Northern-Europe-and-the-Caucasus-scripts.

## Data Availability

Raw sequence data and the *Y. pestis* aligned reads are available through the European Nucleotide Archive under accession no. PRJXXXXXXX.

