## Supplementary Notes for "The Genomic Landscape of Post-Black Death Epidemics in Northern Europe and the Caucasus"

**Contributed equally*

### Supplementary Note 1: Description of Sites and Sampled Individuals

A full list of the genomic coverage and individuals sampled is provided in **Supplementary Table S1**. Here we describe other contextual details of the sites and individuals considered.

#### Lund samples contextual information on the site

Lund, in southwestern Sweden, was during the Middle Ages situated in the prosperous Eastern part of the kingdom of Denmark. Lund was the archiepiscopal center of the Nordic countries with 26 churches and monasteries. It was Denmark's largest mint with flourishing trade and business. However, its importance declined during the fifteenth century as the ports of Malmö and Copenhagen expanded. This decline accelerated following the Reformation of 1536, when the monasteries and most of the churches were demolished.

When Lund was founded c.970-990, it was a political center of power and church. Clergy and the regent's deputies, coiners, artisans, merchants and free and unfree workers of various kinds gathered. The written sources of the city's earliest history are few. But the many archaeological investigations that have been carried out since 1890, in the often well-preserved, thick cultural layers have given a good picture of settlements and everyday life. The surveys also show that the locality did not grow out of a pre-existing village or trading post.

The cemetery of Trinitatis church, where the individuals yielding *Y. pestis* genomes considered in this study were recovered, was the largest medieval cemetery in Lund, and based on excavated areas and the grave density, it is estimated that around 5700 individuals were buried there[1]. The date of the creation of Trinitatis churchyard is based on a dendrochronological dating of one of many wooden coffins. A plank in the coffin came from a tree felled in 985-997, probably during the reign of King Sweyn Forkbeard[2,3]. Based on stratigraphy, dendrochronology and radiocarbon dating in combination with the time-characteristic arm position of the buried individuals, the graves have been divided into chronological groups (990-1050/60, 1050/60-1100, 1100-late 1200, 1300-1536). This latter period was characterised by increased funerary activity in general and by the presence of several double and triple graves. This concentration of graves is interpreted as a reflection of the victims of the Black Death along with subsequent outbreaks in Lund[1].

Our dataset identified 19 individuals positive for *Y. pestis* in Lund Trinitatis cemetery, of which 16 showed sufficient evidence to proceed to genome-wide capture enrichment.

#### Trondheim samples contextual information on the sites

The Norwegian city of Trondheim (medieval Nidaros) was established at latest by 970-980 CE[4]. Founded during the consolidation of the Norwegian kingdom under Harald Fairhair and his successors, the town rapidly developed into one of the most important political and economic centres in Scandinavia.

The foundation of the Archdiocese of Nidaros in 1152/53 established Trondheim as an ecclesiastical capital, overseeing dioceses across mainland Norway, Iceland, Greenland, the Faroe Islands, Orkney, and the Isle of Man. From the 11th century until the Reformation in 1537, Nidaros was the most important pilgrimage destination in Scandinavia, centred on the shrine of St. Olav[5]. This ecclesiastical prominence reinforced the town's demographic and economic growth throughout the High Middle Ages.

The social and demographic impact of the Second Plague Pandemic in Norway is well documented[5,6]. The Black Death reached Norway in 1349 CE, most likely via maritime trade networks, and spread rapidly along the coast. Contemporary and later sources indicate catastrophic mortality, with national population losses often estimated at 50-60%[6].

Trondheim, as one of Norway's largest and most internationally connected towns, was directly affected. Archaeological, documentary, and demographic evidence suggest a dramatic contraction in the population size following 1349, compounded by recurrent plague outbreaks in the later 14th and 15th centuries. The urban population did not recover to pre-1349 levels until the 17th century. This prolonged demographic depression had lasting consequences for settlement structure, land use, ecclesiastical institutions, and burial practices within the town.

The skeletal material analysed in this study derives from two geographically proximate cemetery contexts in central Trondheim, the Library Site and the West Front:

*Library Site*

The so-called "Library Site" cemetery is located just north of the ruins of St. Olav's Church in central Trondheim. Excavations conducted in advance of modern development revealed an extensive medieval burial ground associated with a parish church[7]. Stratigraphic and contextual evidence securely date the cemetery's primary use to before 1349 CE, i.e., prior to the arrival of the Black Death in Trondheim. The burial assemblage is consistent with normative Christian practices of the High Middle Ages: extended supine inhumations, east-west orientation, and limited grave goods. Despite this the individuals included from this context in our dataset (n = 6, four yielding usable *Y. pestis* genomes) likely mostly represent a post-Black Death urban population.

Archaeological and historical evidence indicate that parish churchyards in medieval Norwegian towns generally accepted individuals from across the socioeconomic spectrum, rather than representing a restricted elite population. However, in this case the Library Site is believed to generally represent a higher status segment of the population. The location of the cemetery within the dense urban fabric of medieval Trondheim suggests that it reflects the resident parish population of a thriving town at the height of its ecclesiastical and political significance.

*West Front Cemetery, Nidaros Cathedral (17th-19th Centuries)*

The second site is located at the west front of Nidaros Cathedral. This cemetery area contains burials dated between the 17th and 19th centuries (n = 1 in our dataset), based on stratigraphy, associated material culture, and historical documentation[8].

Following the Reformation in 1537, Trondheim's ecclesiastical structure changed substantially, yet the cathedral remained a central burial location. By the early modern period, burial grounds around the cathedral and associated parish churches continued to receive individuals from a broad cross-section of urban society. Coffin burials, occasional textile preservation, and post-medieval artefacts are typical of this phase. The West Front cemetery thus represents a post-medieval urban population living several centuries after the demographic nadir caused by the Second Plague Pandemic. By the 17th century, Trondheim had begun to re-expand demographically and economically, although it did not reach its estimated pre-1349 population size until this period. In medieval and early modern Trondheim, burial within parish cemeteries was structured primarily by parish affiliation and religious norms rather than strict socioeconomic segregation, although proximity to the church building could reflect status differentiation.

**Armenia sample contextual information on the site**

The site of Aghtsk is located in the Aragatsotn region of the Republic of Armenia, approximately 35 km west of Yerevan, within the central area of the modern village. Archaeological investigations have identified an extensive medieval cemetery surrounding the village church and a royal mausoleum traditionally associated with the Arsacid (Arshakuni) kings, with the funerary complex occupying a prominent position within the settlement[9]. All burials yielding evidence of *Y. pestis* infection were recovered from the upper stratigraphic layer, dated to approximately 1400-1600 CE. These burials predominantly consisted of children or individuals whose mortuary treatment appears to deviate from standard Christian practices, potentially reflecting responses to the cause of death or perceived contagion.

From this context, three individuals yielded *Y. pestis* genomes of sufficient quality for analysis which we detail in turn.

CB010 / Aghck2016_62 was found in a shallow burial (30-50cm top layer) containing at least three individuals represented by disarticulated cranial and long bone remains. The excavators considered the burial atypical of normative Christian funerary practice and suggested that the unusual treatment may reflect exceptional circumstances surrounding death.

CB044 / Aghck2016_57 was an archaeologically assigned male individual recovered in the top layer 30-50 cm from the ground. The corpse was buried according to the Christian rite, lying on its back, arms crossed over his chest. However, due to being on the upper layer, the bones of the skeleton were displaced.

CB045 / Aghck2018_220 was an adult individual aged approximately 30-40 years. The individual was buried in a supine position with the head to the west and face oriented towards the northeast. The arms were crossed over the chest, with the right forearm over the left, and the legs were extended. The burial belonged to the second burial horizon, at a depth of 50-70 cm, adjacent to a wall dated to the fourteenth-fifteenth centuries. The grave was covered by a layer of small to medium sized tuff and basalt stones. The preserved skeleton measured approximately 158 cm in length. The skull was damaged, and the phalanges and metatarsals of the feet were absent.

**Eindhoven sample contextual information on the site**

Eindhoven was newly founded as a marketplace in the early 13th century at the location of a former farmstead and nearby the motte-and-bailey castle Die Haghe. Radiocarbon dates point out that people were buried in Eindhoven from the foundation of the town, but it took until the early 14th century before Eindhoven had its own church, the St. Catherine's church, next to the cemetery[10]. The cemetery went out of use around 1850 CE and until then had served as the only cemetery for the inhabitants of Eindhoven, with the exception of a small Jewish cemetery outside of the town that started in the mid-18th century.

Four samples from Eindhoven yielded detectable *Y. pestis* DNA, with one individual taken forward for capture enrichment and considered in this study due to its coverage (EIN2927), **Supplementary Table S1**. EIN2927 was part of a wider collection recovered during a single excavation that took place between March 2005 and August 2006[11]. The excavation consisted of one trench covering the eastern extensions of the former late medieval/Early Modern Period Catherina's church and part of its northern, eastern and southern cemetery and reached the undisturbed soil. Altogether, this means that people from throughout the entire period of use of the cemetery and different social-economic layers within the community were excavated.

EIN2927 (S3489, M2927) was located in the southern part of the cemetery and was found at the edge of the excavation trench. During the excavation, no mass burials were found and there is no specific mention of plague cemeteries in Eindhoven, though it is known that plague victims were buried at the Catharina cemetery where this individual was found. EIN2927 is a female individual of age 40-49[12]. EIN2927 was found underneath a burial (feature S2196) with a 14C date of 1454-1632 calCE (GrA38363, 370 +/- 25 BP)[12]. Based on archaeological context, we conservatively estimate the archaeological date of this individual to 1200-1632.

**Icelandic sample contextual information on the site**

USK13 is a female individual recovered from a site called Útskálakirkjugarður (Útskála cemetery) in Reykjanes Peninsula. The part of the cemetery from where this individual was excavated is thought to have been in use during the 13th-17th centuries. The churchyard itself was in use for centuries, with the oldest standing headstone dating back to around 1600. Hundreds of disarticulated human skeletal remains were recovered from under the floor of the church during construction work in 2007, which included the tooth sample considered here; a mandibular second premolar. All bones were fully disarticulated and mixed, so there is no individual burial context available for USK13, making the site poorly dated from an archaeological standpoint. The radiocarbon age of this sample is 411 ± 27 BP (SUERC-105842), calibrating to 1443-1485 calAD (68.3%) on IntCal20 without marine correction. The isotope values (δ13C = -18.7‰, δ15N = 12.0‰) indicate a significant marine dietary signal, hence recalibration was conducted using a mixed marine/terrestrial curve in OxCal 4.4 specifying 27.1% marine (linear interpolation following Arneborg *et al.* 1999[13]) and a local Icelandic reservoir correction of ΔR = -76 ± 70 years from the Calib Marine database. The corrected dates point to a date range of c.1460-1814 (68.3%: 1519-1598 CE and 1615-1686 CE and 95.4%: 1460-1814 CE).

### <u>Supplementary Note 2:</u> Sample Processing, DNA Extraction, Library Preparation, Capture Enrichment and Radiocarbon Dating

The majority of the uncaptured data presented in this work comes from an ancient DNA dataset generated from human remains excavated in Lund (Sweden) and Trondheim (Norway)[14]. Supplementing this was unpublished data derived from skeletal material excavated in Eindhoven (Netherlands), Aghtsk (Armenia) and Útskálakirkjugarður (Iceland) (**Supplementary Table S1 and Supplementary Note 1**), as well as data generated specifically for this study from two of the Norwegian samples (SK111 and SK025). Teeth samples were processed in dedicated ancient DNA laboratories at the Globe Institute, University of Copenhagen, Denmark. DNA was extracted and sequenced from unpublished samples as follows:

Before sampling, the teeth were cleaned by wiping them with a 1% sodium hypochlorite solution. Teeth were sectioned at the cementoenamel junction using a diamond blade, and 36-51 mg powder was drilled from inside the pulp chamber of the molar crown. Note for samples published in Liu *et al.* the cementum was instead targeted due to the focus of this study on recovering human DNA[14]. DNA was extracted from the tooth powder following a modified version of Moreno-Mayar *et al*. 2018's[16] protocol, 1.2 ml of digestion buffer was added to each sample, after which they were incubated overnight at 37°C on a rotor. The digestion buffer consisted of 0.463M EDTA pH8, 10mM Tris, 0.5% N-laurylsarcosine, 30%, 1x Phenol red and ca 0.2 mg/ml Proteinase K. DNA was purified from 1 ml of the digest using a modified version of Qiagen PB buffer (cat: 628 19066) and centrifugation through Roche large volume silica columns (cat: 05114403001) according to McColl *et al*. 2025[17]. The DNA was eluted using 65µL Qiagen EBT buffer into 1.5ml Eppendorf LoBind tubes. DNA extracts were then converted into single-strand DNA Illumina libraries using the Santa Cruz Reaction (SCR) protocol of Kapp *et al*. 2021[18].

While most of the sequence data derive from full-UDG treated libraries used for genomes reconstruction (see *Y. pestis* whole genome capture), libraries for most of the samples were initially prepared without UDG treatment to assess DNA damage profiles along with the screening data which did not undergo UDG treatment (**Supplementary Table S2**). Prior to indexing, we performed quantitative real-time PCR (qPCR) with SYBR Green to determine the optimal number of amplification cycles per sample. The libraries were cleaned using SPRI beads at a ratio of 1.2X to remove adapter dimers, followed by quality control on a Bioanalyzer. Samples were pooled at equimolar concentration in batches for deep sequencing.

#### Screening with MALT and HOPS

All non-UDG treated shotgun sequenced DNA from Trondheim and Lund (see Liu *et al.*[14]) and new libraries from Trondheim, Eindhoven, Icelandic and Armenian sites were screened for traces of ancient pathogen DNA, totalling approximately 660 individuals screened (**Supplementary Table S3**). The raw fastq files were pre-processed using the nf-core/eager[19] pipeline (v2.3.5). fastp[20] (v0.20.1) was used for poly-G tail removal. Subsequently, AdapterRemoval[21] (v2.3.1) was used to remove adapter sequences and to merge paired-end data. The adapter-clipped and merged fastq files were analysed with MALT[22] (v0.4.1) using a custom database constructed using complete bacterial, viral and archaeal genomes as previously described[23]. The --malt-run command was executed in BlastN mode with SemiGlobal alignment. The --minPercentIdentity parameter was set to 95, --minSupport was set to 10 and --topPercent to 1. Remaining parameters were set to default. The MALT output rma6 files were subsequently screened for pathogens using the MaltExtract and AMPS tools integrated in the HOPS[24] pipeline. A previously published custom taxon list[25] (github.com/ashildv/custom_hops_pathogen_list_v4) was used to screen for pathogens with HOPS. Additionally, all rma6 MALT output files were visually inspected using MEGAN6[26].

#### *Y. pestis* whole genome capture

Based on the pathogen screening results, 32 individuals were chosen for *Y. pestis* whole genome capture (**Supplementary Table S1**). Five individuals - SK051 (Trondheim), LUN375, LUN402, LUN431 (Lund), EIN5221 (Eindhoven) - were excluded from whole *Y. pestis* genome capture as they were deemed to be too weakly positive for *Y. pestis* and unlikely to yield a meaningful amount of reads from genome capture.

New subsampling of the teeth and subsequent DNA extraction was carried out for 29 of the individuals to generate sufficient genetic material for whole genome capture. In all cases 17-76 mg of powder was drilled from inside the tooth crown pulp chamber. For four individuals (CB045, CB048, CB073, LUN105) entirely new teeth were sampled, and in two cases (CB010, SK110) two teeth were sampled for the same individual, in both cases one was the tooth already sampled and the other was a new tooth. DNA was extracted from the newly sampled powder as described above for tooth powder samples and converted into UDG treated single stranded libraries by incubating the extracted DNA with Thermolabile USER® II Enzyme (NEB) for 3h at 37℃ followed by 10 minutes of enzyme heat inactivation at 65℃ prior to using the SCR protocol[18]. A full overview of the sampling, extracts and libraries generated for capture is presented in **Supplementary Tables S1-2**.

All individuals/samples were assigned a separate capture pool. The recommended amount of input DNA when capturing ancient DNA samples using RNA bait hybridization capture with myBaits® (Daicel Arbor Biosciences) is 1 µg or more. For all individuals from Trondheim, Eindhoven, Lund and Armenia, 95 % of the DNA amount per sample pool was made up of UDG-treated library and the other 5 % of the corresponding non-UDG library. In five cases (SK079, LUN099, LUN436, EIN2082, EIN1788), the DNA pools submitted for capture were below 1 µg, ranging from 633-836 ng. For the Icelandic individual USK13, the DNA pool was made up of two non-UDG-treated libraries. All indexed negative control libraries were combined at approximately equimolar concentration into one capture pool. In total, 35 capture pools were prepared.

All 35 pools were shipped to Daicel Arbor Biosciences' location in Ann Arbor, Michigan, U.S.A on dry ice, where the *Y. pestis* whole genome hybridization capture was performed by trained professionals using RNA baits/probes. The capture was done using a myBaits®180-200k kit. A previously designed probe set published by Wagner *et al.* 2014[27] was used to capture for the *Y. pestis* genome and associated plasmids, with one alteration to the pPCP1 probes. The region of the pPCP1 genome spanning positions 3000-4200 (NC_003132.1) was excluded from the probe design. This region was removed because it is similar to an expression vector used in enzyme production and is known to capture off-target DNA fragments, as was demonstrated by Schuenemann *et al.* 2011[28]. In brief, the probe design consisted of baits for the core *Y. pestis* chromosome (+pMT1 & pCD1 plasmids) designed across a range of *Yersinia* genomes. The design also includes a set of additional probes to increase coverage of SNPs of phylogenetic interest/importance, plus a low-level spike-in of probes for plasmid pPCP1. The probes in this design are 80 bp long with 20 bp tiling density.

### Radiocarbon dating

18 samples for which there was sufficient material remaining were sent for radiocarbon dating at the facilities at NTNU University Museum in Trondheim, Norway (https://www.ntnu.edu/museum/services); as described in Liu *et al*. 2026[14]. Radiocarbon dates were calibrated in OxCal v4.4 using the IntCal20 terrestrial and Marine20 marine calibration curves. For individuals with evidence of marine dietary input (discussed in the next section), mixed-source calibrations were performed using estimated marine dietary proportions derived from $\delta^{13}C$ values, together with regional ΔR corrections. Uncertainty in dietary estimates was incorporated by allowing at least a ±5% error around the estimated marine contribution, or more when estimated marine dietary proportions derived from $\delta^{13}C$ and $\delta^{1}5N$ differed substantially. The methodology is described in Seiler *et al*. 2025[29], based on the Medieval material from Trondheim. For samples from Lund we know the cemetery was in use until 1536, so throughout this was set as an upper bound on the dates. An overview of the radiocarbon dating results is presented in **Supplementary Table S1**.

### Assessment of Marine Diet

For partial marine diets, the radiocarbon dates needed to be corrected for an according proportion of the marine reservoir effect. This is relevant for all regions and periods where the consumption of marine resources was an important contribution to the diet and especially important during the Medieval period, where fasting rules prohibited the consumption of meat on about half of the days in a year, and where corrections of a few decades can mean significant changes of the interpretation of the data.

We calculated both $\%marine_{\delta 13C}$ and $\%marine_{\delta 15N}$. A comparison of the two %marine values from each individual was then used as a test to check for internal consistency. For the reservoir corrections, we used the %marine as calculated from $\delta^{13}C$, because that is (in absence of C4-based diet) a more direct reflection of marine food intake. $\delta^{15}N$ is correlated with %marine as well, but can also increase due to other factors such as consumption of high-trophic level terrestrial food, nursing, pregnancy, starvation or water stress. The uncertainty of the calculated %marine was calculated as the difference between $\%marine_{\delta 13C}$ and $\%marine_{\delta 15N}$, or at least 5% when that difference was below 5%.

The skeletons analysed here were from different geographical areas with different isotopic baselines, thus needing different reservoir corrections. We assumed that a regional correction is the most appropriate, knowing that mobility may blur the picture (the isotopes in a person's tooth reflect the diet of the time when that tooth formed, which could have happened in another region than the place of burial).

For the samples from Armenia, we suspected no significant marine contribution. The high $\delta^{13}C$ values are probably due to the consumption of a diet based on plants with C4 photosynthesis, e.g. millet (including animals that consumed C4 plants such as certain grasses). The samples from Lund and Trondheim were both from Scandinavia, but still geographically different:

Trondheim is in central Norway at the North Sea coast, while Lund is in southern Sweden at the coast of the Øresund (that connects the Baltic to the North Sea).

For the samples from Lund, we used similar baseline values as those found for Medieval Copenhagen[30], assuming that the geographical closeness of Lund and Copenhagen across the Øresund is reflected in the isotopic baseline. Some differences were observed, though. The human bones from Copenhagen had $\delta^{13}C$ values between -19.7 and -17.3 ‰, $\delta^{15}N$ between 10.0 and 15.5 ‰. This is higher, but overlaps with the ranges we found for Lund, $\delta^{13}C$ between -20.2 and -18.3 ‰ and $\delta^{15}N$ between 8.8 and 14.4 ‰. Olsen *et al.* 2019 used $\delta^{13}C$ between -20 ‰ (terrestrial) and -10 ‰ (marine). As some of the individuals from Lund had values below -20 ‰ (-20.03±0.03, -20.18±0.08 and -20.15±0.06), our baseline needed to be slightly lower. For the terrestrial endpoint, we thus used -21 ‰. For the marine endpoint, we chose to use -12 ‰ instead of the -10 ‰ suggested by Olsen *et al.* 2019 - not a result of direct measurements on marine resources, but considering that the %marine diet reconstructions based on $\delta^{13}C$ and $\delta^{15}N$ values generally should agree. For the terrestrial $\delta^{15}N$ endpoint we used 8.5 ‰, as the human bone in our assemblage with the lowest value had a value of 8.84 ‰. This is significantly lower than those found for e.g. Trondheim[29], where a terrestrial baseline of $\delta^{15}N$=10 ‰ was found to be the best representation of the population's isotope values. Variations in the terrestrial $\delta^{15}N$ values can be caused by factors such as climate, land use history, and/or agricultural practices such as manuring. For the marine $\delta^{15}N$ endpoint, we used $\delta^{15}N$=23 ‰, based on the most extreme values found in the literature and measured for human bones throughout the laboratory's history (National Laboratory for Age Determination, NTNU University Museum, Norwegian University of Science and Technology).

The correlation between $\delta^{13}C$ and $\delta^{15}N$ was weak for the Lund individuals ($R^2$=0.26). This was probably caused by a few individuals who have higher $\delta^{15}N$ values than predicted by marine diet. These could have had a higher trophic level diet, indicating higher social status, or the elevated $\delta^{15}N$ values could have been caused by breastfeeding, starvation, disease or pregnancy. LUN409, but also LUN112 and LUND099 ,seem to lie on another trajectory than the rest.

For the Trondheim sample (TRa-20048, SK110), we used the isotopic baselines developed on the basis of a study of a large skeletal assemblage from several cemeteries from this town[29]: $\delta^{13}C$=-22 to -12 ‰ and $\delta^{15}N$=10 to 23 ‰.

### Supplementary Note 3: Processing Ancient and Contemporary Genomic Data, Quality Control and Reproducible SNP Filtering

Raw sequencing data were processed using the nf-core eager pipeline[19] (v2.4.7) implemented in Nextflow[31] (v22.10.0). Adapters were removed using AdapterRemoval[21] (v2.3.2) with the minimum adapter overlap set to 3; additionally, samples that were sequenced with a two-colour chemistry had poly-G tails removed using fastp[20] (v0.20.1). Full-UDG-treated libraries were mapped using BWA aln[32] (v0.7.17) with stringent conditions (-n 0.1, -l 32). Following the mapping, the bam files were filtered using samtools[33] (v1.12), requiring a mapping quality of 37; unmapped reads were discarded, and deduplication was performed using Picard MarkDuplicates[34] (v2.26.0).

Half-UDG treated libraries were mapped using BWA aln with lenient conditions (-n 0.01, -l 16). Following the mapping, the bam files were filtered using samtools, requiring a mapping quality of 37, and unmapped reads were discarded. After this initial mapping, the reads were extracted from the bam files using samtools fastq and the first and last base pair of the reads were trimmed using fastp. Following this, the reads were remapped using BWA aln with stringent conditions (n 0.1, -l 32). The bam files were filtered using samtools with a mapping quality of 37, unmapped reads were discarded, and deduplication was performed using Picard MarkDuplicates.

Non-UDG treated libraries were separated into libraries which could be merged with libraries with other types of UDG treatment and those that could not. Non-UDG libraries that could not be merged with libraries with other kinds of UDG treatments were mapped using BWA aln with lenient conditions (n 0.01, -l 16) and underwent Bayesian rescaling using mapDamage2[35] (v2.2.1) specifying a sequence length of 30. Following the mapping, the bam files were filtered using samtools, requiring a mapping quality of 37, and unmapped reads were discarded. After this initial mapping, the reads were extracted from the bam files using samtools fastq and remapped using BWA aln with stringent conditions (n 0.1, -l 32)[36]. Following the mapping, the bam files were filtered using samtools with a mapping quality of 37, unmapped reads were discarded, and deduplication was performed using Picard MarkDuplicates.

Non-UDG libraries merged with libraries with other types of UDG treatments were mapped using BWA aln with lenient conditions (n 0.01, -l 16). bamUtil[37] (v1.0.15) was used to clip the first and last five base pairs of the reads. Following the mapping, the bam files were filtered using samtools with a mapping quality of 37, and unmapped reads were discarded. After this initial mapping, the reads were extracted from the bam files using samtools fastq and remapped using BWA aln with stringent conditions (n 0.1, -l 32). Following the mapping, the bam files were filtered using samtools with a mapping quality of 37, unmapped reads were discarded, and deduplication was performed using Picard MarkDuplicates.

Where necessary, the resulting library bam files were merged using samtools merge, deduplication was performed using Picard MarkDuplicates and the bam file was then sorted and indexed using samtools.

#### Processing modern *Y. pestis* genomic data

Where possible, raw sequencing data were downloaded for each sample. When this was not possible, the assembly was downloaded and simulated reads were generated using pyfasta (0.5.2). The assemblies were split into 100-mers with 99 bp overlap, and the fasta files were converted to fastq files with artificial base quality scores using the fasta_to_fastq perl script (https://code.google.com/archive/p/fasta-to-fastq/). Sequencing data was processed using the nf-core eager pipeline implemented in Nextflow[19,31]. Adapters were removed using AdapterRemoval with the minimum adapter overlap set to 3. Libraries were mapped using BWA mem with default parameters. Following the mapping, the bam files were filtered using samtools, requiring a mapping quality of 37; unmapped reads were discarded, and deduplication was performed using Picard MarkDuplicates.

**Genotyping and SNP filtering**

The filtered and deduplicated bam files across modern and ancient strains with a mean coverage greater than 5x were processed using the nf-core eager pipeline. Genotyping was performed using GATK Unified Genotyper[38] (v3.5.0) with the EMIT_ALL_SITES flag. SNP tables and alignments were generated using MultiVCFAnalyzer[39] (v0.85.2), requiring a minimum genotyping quality of 30, minimum base coverage of 3x and a minimum allele frequency of 0.9 for homozygous calls. The genome of *Y. pseudotuberculosis* IP32953 was included as an outgroup, and regions previously identified as non-core regions containing repetitive elements or coding for tRNAs, rRNAs or tmRNAs were masked[40]. The SNP table and SNP alignment output by MultiVCFAnalyzer were imported into Julia using the CSV.jl, DataFrames.jl and FastaIO.jl (1.1.0) packages. Coverage information on each of the genomes was generated using samtools and used to construct a coverage matrix.

A series of SNP filtering steps was then applied. Firstly, sites identified as multiallelic were excluded from the SNP alignment. Singleton sites were identified and subsequently excluded if there was another singleton within 50 bp either side of the singleton or where coverage dropped to 0 within a 100 bp window around the singleton. SNPs, where calls were missing with a frequency of more than 0.23 in ancient genomes, were also excluded. Further, any singleton with a coverage that was more than three standard deviations above or below the alignment's mean coverage or suspected of being an erroneous call due to deamination damage was also excluded. In addition, the dataset was interrogated for any reported singletons in genes (*recA* and *mutS*) associated with hypermutator phenotypes.

Singletons were considered as potentially deriving from post-mortem damage if they occurred in an ancient genome and exhibited a base change typically attributed to aDNA (G->A, C->T). The coverage for all SNPs in the genomes that contained potentially erroneous calls was extracted, and the mean and standard deviation were calculated. This information was then used to calculate a z-score for the coverage of the positions where the potentially erroneous calls were made. The minimum coverage threshold for accepting the call was set to 5-fold or a z-score of -1, whichever was stricter. Calls that did not meet this stricter threshold were excluded from the SNP alignment.

The resulting alignment, comprising 2834 SNPs, was used to generate a phylogenetic tree (see **Phylogenetic reconstruction**). This tree, along with the masked SNP alignment, was used to identify homoplastic sites using HomoplasyFinder[41]. The resulting consistency index report was used to identify homoplastic sites. Sites identified as homoplastic, indicated by a Consistency Index of less than 0.5, were excluded from the alignment. The phylogenetic tree was then reconstructed using the resulting alignment of 2804 SNPs, and a matrix of nucleotide differences was generated using snp-dists[42] (v0.8.2). While this number is lower than some previous phylogenetic studies of *Y. pestis*, we took an ultra-conservative approach to ensure a robust final topology. The final SNP calls are provided in **Supplementary Table S6**.

**Identification of genomes for exclusion from downstream analysis**

While conducting our SNP filtering analysis, we observed that some of the included *Y. pestis* genomes SQ8723_SQ8833_SQ8962, LAR27 and AMIENS268 sat on unusually long terminal branches. To further investigate this, we began by extracting and plotting mutation positions across the genomes using Makie.jl in Julia, finding these tended to be clustered (**Supplementary Figure 1**). Observing, in particular, an unusual peak of mutations shared between these three genomes at 210-240kb, we extracted all reads that overlapped with these mutations and used NCBI blast-n to determine whether they likely derived from possible contamination with non-*Y. pestis* species. On finding these were mostly non-*Yersinia* in origin (**Supplementary Figure 2-4**), we decided to exclude these genomes from further analysis.

### **Supplementary Note 4:** Bayesian phylogenetic dating

**Bayesian phylogenetic tip dating with Beast2**

The phylogenetic tree and associated tip dates were imported into TempEst (v1.5.3) and the root-to-tip correlation was assessed to confirm the presence of a temporal signal[43] (**Supplementary Figure 8**). Following the methods in Spyrou *et al.* (2022), the following model parameters were implemented in BEAUti (v2.7.8): the site model was set as a General Time Reversible (GTR) model with four gamma rate categories, and the AG substitution rate parameter was fixed at 1.0. The clock was configured to an optimised relaxed clock rate with a uniform prior distribution ranging between 1e^-3 and 1e^-6 substitutions per site per year for the SNP alignment. The tree prior was set to a coalescent skyline, and a Jeffreys prior distribution was utilised for the population sizes. A dimension of five was employed to allow for variations in group and population sizes over time, with an upper limit of 380,000 for the effective population size. Additionally, all Branch 1-4 genomes (both ancient and modern), as well as the 0.ANT3 lineage were constrained to be independent monophyletic clades.

This analysis was conducted in BEAST2[44] (v2.7.7) using three independent chains, which were then combined using LogCombiner (v2.7.7). Convergence was assessed using Tracer[45] (v1.7.2), ensuring that the effective sample sizes (ESS) exceeded 200 for each estimated posterior distribution after a 10% burn-in period. Maximum clade credibility trees were constructed using TreeAnnotator (v2.7.7) within BEAST2, also applying a 10% burn-in, and were subsequently visualised in R using the ggtree package.

**Bayesian phylogenetic tip dating with BactDating**

A maximum likelihood tree of branch 1 genomes was generated with IQTree (see **Phylogenetic Reconstruction**). The phylogenetic tree and associated tip dates were imported into TempEst (v1.5.3) and the root-to-tip correlation was assessed to confirm temporal signal[43] (**Supplementary Figure 8**). The resulting tree was imported into R with the package ape (v5.8)[46], and the tree was rooted to the outgroup (*Y. pseudotuberculosis IP32953*) and the outgroup was dropped from the tree. The branch lengths were then converted from substitutions per site to units of substitutions. Following this, polytomies were removed from the tree with the multi2di function which works via transforming all multichotomies into a series of dichotomies with one (or several) branch(es) of length zero. After this initial set up, the tree was read into BactDating (v1.1.2)[47] along with date ranges capturing the upper and lower bounds of the uncertainty around the sample dates. The strict gamma model was used along with a coalescent tree prior and the update root flag was set to false. After $5e^{+6}$ Markov chain Monte Carlo (MCMC) iterations were performed, the ESS and traces of the parameters were interrogated to ensure MCMC convergence. The tree was then visualised with the ggtree package (**Supplementary Figures 8-10**).

**<u>Supplementary Note 5:</u>** Phylogenetic association of *Y. pestis* genomes considered in this study with historically attested outbreaks
*Philip Slavin, University of Stirling*

To link our newly generated *Y. pestis* observations to historically recorded outbreaks we contextualised our findings aided by the framework developed by Keller *et al*. (2026)[48]. In this work they employed PhIRM (phylogenetically informed radiocarbon modelling) to improve the dating of plague burials by incorporating phylogenetic data into Bayesian radiocarbon modelling. They subsequently compared the resulting chronological ranges with documentary evidence, thereby connecting the sequenced and examined genomes with a set of documented plague outbreaks (typically between one and five). Each previously published genome incorporated into Keller *et al.* (2026) was extensively contextualised and matched with probable historically attested outbreaks. This serves as the foundation for linking the newly sequenced and analysed genomes presented in this study with their corresponding historical outbreaks, based on their phylogenetic placement relative to the genomes published, analysed, and reassessed in Keller *et al*. (2026).

**Supplementary Table S8** gives an overview of the genomes included in this study as an ancient reference dataset. On the basis of (1) Keller *et al*. (2026)'s proposed associations with historical outbreaks and (2) the phylogenetic positioning of the genomes examined in the present study relative to those analysed by Keller *et al.* (2026), we associate the genomes from this study with specific historical outbreaks in turn.

The use of branch lengths to determine the number of SNPs partially accounts for missingness in the sites covered but it remains that SNPs on terminal branches may not be well typed. As such, the SNP differences discussed here, and resulting assignments, should be taken as a minimum number, and results, particularly on placed lower-coverage genomes where terminal branch lengths are not biologically meaningful and we rely on a mixture of internal branch lengths and the SNP distance matrix (**Supplementary Table S11**), should be considered with appropriate caution.

#### Lund *LUN105, LUN106, LUN 124 and LUN252* genomes

The Lund genomes LUN105, LUN106, LUN124 and LUN252 are positioned together with other known Black Death genomes, implying they, too, are to be associated with the first wave of the Second Pandemic. According to textual evidence, plague appears to have arrived in the Västergötland region of Sweden from southern Norway in late 1349, and spread all over southern Sweden, including Lund, in 1350[49–51]. The 1350 mortality in Lund is reflected in the increased number of individuals' deaths mentioned in a local Cathedral obituary book, while in the autumn of the same year, there was an increased number of pious donations to the Cathedral[52].

#### *Pestis secunda* genomes: Lund *LUN132*, Aghtsk *CB010*, *CB044* and *CB045*

The Lund *LUN132*, Aghtsk *CB010*, *CB044* and *CB045* genomes are all positioned on Branch 1B and, therefore, are associated with the *pestis secunda* wave, commencing most likely in

South-Central Germany in summer 1356 and ravaging all over Europe, the Middle East and North Africa in 1356-1366[53]. During the same wave, plague was documented in Lund in 1359, whereby excess mortality was reported in a local necrolog[54]. Likewise, plague was reported in other regions of Sweden, including Gotland and Uppsala, in the same year[54].

It was not until 1364 that the *pestis secunda* reached the South Caucasus, having likely arrived from the Pontus, Crimea, the Pontic–Caspian Steppe, North Caucasus, or a combination of these regions[53,55]. In Armenia, a plague outbreak is mentioned, in general terms, in 1364-1365[56]. Importantly, CB045 is positioned together with the Bolgar Fort genome (Bolgar2370), implying that both Bolgar and Aghtsk may have experienced infections with a strain from the same source, situated somewhere halfway in-between: that is, somewhere in the Pontus, Crimea, the Pontic-Caspian Steppe or North Caucasus.

**Lund *LUN214, LUN363U* and Útskálakirkjugarður *USK13* genomes**

At some point between the Black Death and the *pestis secunda* waves (that is, between c.1348 and 1356), Branch 1 (responsible for the Black Death waves) underwent a split into the 1A and 1B sub-branches, with the latter causing the *pestis secunda,* and the former triggering later 14th- to 19th-century outbreaks in Europe. Shortly after the 1A-1B split, there was another split within the 1A branch, trifurcating into three sub-branches, with one clustering the OTE001 (Otepää, associated by Keller *et al.* 2026 with one of the following outbreaks: c.1368, 1378-1379, c.1389-1390) and COL1 (Collalto Sabino, associated with either the 1374 or 1384 outbreak) genomes; the other leading to a further trifurcation into the branches represented by (1) MAN008 (Manching-Pichl, associated with one of the following outbreaks: 1371, 1373, 1380, 1396), (2) X3003 (Sejet, which could be associated with any of the following outbreaks: 1495-1496, 1511, 1520, 1527, 1546-1548, 1553-1554, 1563-1568, 1575-1578, 1583-1585, 1592-1594, 1601-1603, 1618-1621, 1624-1626, 1629-1630, 1636-1638) and (3) LUN214/363U (Lund) and USK13 (Útskálakirkjugarður) genomes; and the third one acting as a 'main' 1A branch leading to later nodes birthing new sub-branches and causing subsequent plague waves. It is possible that the trifurcation is associated with the *pestis tertia* commencing in South Germany (possibly in Franconia) in 1362, or the subsequent wave commencing, in a back-to-back manner in two South-German regions (possibly in the Bodensee/Appenzell and Franconia, respectively) in 1377 and 1379. If this is true, then we may date the event to c.1362-c.1379.

There are five SNPs between the trifurcation and the LUN214/363U genomes. Despite some attempts to calculate 'average' SNP accumulation rates of *Y. pestis*[55,57–59], these were anything but fixed and constant, varying not only across branches, but also within the same branches, experiencing phases of deceleration and acceleration[48,60,61]. All the same, the rate of one mutation every 4.8 or so years, estimated for the pre-Little Polytomy (the 1A1-1A2 split c.1450-1500) phase by Keller *et al.* 2026 may still be used as a very approximate guideline here. Multiplying 5 SNPs by the factor of 4.8 renders about 21 years separating the trifurcation and the LUN214/363 genomes. As noted above, Keller *et al.* 2026 associated MAN008 with one of the following outbreaks reported in Bavaria: 1371, 1373, 1380, 1396 (yielding the approximate chronological range of c.1371-c.1396). Hence, the dating of the LUN214/363U genomes may be very roughly approximated to c.1392-c.1417 range, which is partly consistent with their radiocarbon dating to c.1393-c.1460. Within the c.1392-c.1417 range, there were documented and inferred outbreaks in Lund and other regions in southern Sweden in 1387-1389, 1405 and 1413[54], and theoretically the LUN214/363U genomes may be associated with any of these. However, given the wider radiocarbon bracketing of LUN214/363U (c.1393-c.1460), there is always a possibility that they may have been associated with one of later 15th-century outbreaks, discussed in the following sections.

The USK13 genome is nine SNPs derived from the LUN214/363U ones, although this number is not secure, because of the low coverage of the former genome. In any event, it is clear that USK13 is significantly more derived than LUN214/363U and hence, they could not be associated with the same outbreak. As far as plague in Iceland is concerned, there were only two documented outbreaks, both in the 15th century: in 1402-1404 and 1494-1495 - in addition to a number of other epidemic diseases[62,63] that clearly differ in both symptoms and terminology from the plague outbreaks (specifically, the Old Icelandic term for 'plague' is *plága*, as opposed to the more generic *sótt* or *bóla*, meaning, roughly, 'pestilence' or 'epidemic'). Assuming that (1) 9 SNPs are all true, noting we cannot account for missing or private SNPs due to low coverage; (2) the accumulation rate was indeed once in about 4.8 years; and (3) the LUN214/363U genomes overlap in corrected radiocarbon dates to 1393 to 1460, then USK13 may hypothetically fall within the time bracket of c.1436-c.1503 (9x4.8=43.2 years after c.1393-c.1460). This estimate is more consistent with the latter outbreak. There were however other infectious diseases reported in Iceland within this period that could potentially have been caused by plague. In 1426, there was a 'krefðuvetur almikill' ('very severe winter outbreak'), manifesting in eye- and kidney pains, spreading to the chest and causing sores and swelling on the throat and face[64]. In 1430-1431, there was a 'bóla mikil' ('great pestilence') and 'dauði mikill' ('great mortality')[64,65]. In 1462-1463 and 1472 Iceland was again visited by 'mikil bóla'[65,66]. A further epidemic (sótt) was mentioned in either 1479 or 1484, causing the death of a wealthy woman[65].

Unfortunately, USK13 (archaeological ID USK-1-3) was uncovered during the 2007 rescue excavations associated with construction works at Útskálakirkjugarður, with the remains being completely disarticulated and lacking any archaeo-historical context suitable for analysis. The C14 dating rendered the following dating brackets after correction for marine reservoir effects: 68.3%: 1519-1598 CE and 1615-1686 CE; 95.4%: 1460-1814 CE, with the latter covering the 1494-1495 outbreak.

**Trondheim *SK110*, Lund *LUN194, LUN369U*, *LUN112* and *LUN099* genomes**

Following the 1360s/1370s trifurcation, discussed in the previous section, the main 1A branch underwent yet another trifurcation event around c.1398-c.1421, as may be inferred from the dating of the AHM011 (Arnhem) genome, associated by Keller *et al.* 2026 with one of the following outbreaks: 1398, 1410-1412 and 1421, and positioned right on the trifurcation node (implying that the geographic source of the same trifurcation was probably not far from Arnhem). In the course of the event, three branches emerged: (1) one branch, represented by the SK110 genome from Trondheim (which may have been a short-lived terminal branch, although the current lack of other genomes cannot be taken as firm evidence of that); (2) Branch 1A4, represented by published genomes from Denmark, Estonia and newly sequenced genomes from Lund; and (3) the third branch acting as a continuation of the 'main' 1A branch leading to later nodes birthing new sub-branches and causing subsequent plague waves. Branch 1A4 has two subclades, one with the identical genomes of LUN194 and LUN369U, and the other with the genomes A1480x1480 (Denmark) and LUN112, identical with the node giving rise to two terminal branches leading to MAL003 (Mäletjärve, Estonia) and LUN099, respectively.

Trondheim SK110 is a placed low-coverage genome, and hence any estimate of its terminal branch should be considered with appropriate uncertainty. It appears that it is placed at least 8 SNPs derived from AHM011 (c.1398, 1410-1412 or 1421). This potentially places SK110 within the approximate chronological range of c.1430-c.1470. Unfortunately, our information about 15th-century outbreaks in Norway is very limited, but there is a reference to two undated plague outbreaks (most likely referring to, respectively, the years 1430 and 1439, the former generally in Norway and the latter in Bergen), and two further outbreaks in 1452 and 1459, both in Bergen[67–69]. The c.1430 outbreak was also documented in Helsingborg (Sweden) in 1430-1431[70]. The c.1439 outbreak was a part of a much wider trans-European outbreak, ravaging all over Scandinavia in 1439-1441; no further outbreaks were reported in the entire region in the 1440s. The 1452 outbreak was associated with a wave reported all over Denmark and Sweden in 1451-1452 (having been imported from northern Germany)[54,71]. The 1459 outbreak in Bergen was apparently associated with the same wave reported in Denmark in 1460[71].

LUN194/LUN369U are 3 SNPs derived of AHM011 (c.1398-c.1421) - sharing one SNP with the sub-branch, leading to LUN112 and A1480x1480 and derived strains. Hence, LUN194/LUN369U are more derived than AHM011, possibly associated with a subsequent wave. With all the caveats, we may date LUN194/LUN369U, approximately to some ~8-15 years after AHM011, namely to c.1406-c.1436. Within this interval, plague outbreaks were recorded in Sweden in 1404-1405, 1413, 1421-1422, 1430-1431[49,54,70,72]. None of the sources mention an outbreak specifically in Lund. However, the absence of documented outbreaks in Lund in other years does not imply that the city was free of plague; rather, it reflects the lack of surviving textual evidence for those years. Hence, the identical LUN194/369 genomes may be associated with any of these outbreaks.

LUN112 is 5 SNPs derived from AHM011 (c.1398-c.1421) and placed on a node following the bifurcation node between the LUN194/LUN369 and LUND112/LUN009 sub-branches (see the previous paragraph); thus, LUN112 does not have any private SNPs. Roughly speaking, LUN112 may have been some 15-20 years younger than AHM011, and thus hypothetically dated to c.1413-c.1441. Importantly, according to Keller et al. 2026, A1480x1480 is situated close to the Estonia genome MAL003, with the latter being either directly ancestral to A1480x1480 or carrying three private SNPs interpreted as potential damage. In any event, Keller et al. 2026 dated MAL003 to the c.1404-1425 range (listing c. 1404, 1420-1421, 1424-1425 as potentially associated outbreaks), on the basis of the presence of penny of Damerow, or bracteate, minted between 1379 and 1420, found in the same burial (allowing a few extra years for the bracteate to remain in circulation). Moreover, according to the same paper, PhIRM analysis places A1480x1480 within the 1383-1422 range. All this potentially narrows down the dating of A1480x1480 and LUN112 to c.1413-c.1422. Within the said chronological bracket, we hear about plague outbreaks in Denmark in 1420, as well as 1413-1414 and 1421-1422 in Sweden (mentioned and referenced in the previous paragraph)[54,71,72]. It is possible that there was an undocumented outbreak in Denmark in the earlier 1410s, associated with the same wave causing the 1413-1414 outbreak in Sweden. In other words, LUN112 may be associated with either 1413-1414 or 1421-1422 outbreaks.

LUN099 is two SNPs derived from LUN112, forming a short terminal sub-branch. One possibility is that it could have been associated with a later outbreak. Another possibility is that both genomes are associated with the same outbreaks, with LUN099 acquiring two private SNPs in the course of a single local but persistent outbreak. Such variation is not unprecedented, as *Y. pestis* genomes recovered from the same spatio-temporal context have been shown to differ by a single SNP. Examples include Krakauer Berg in Saxony-Anhalt (c.1357), where two of five genomes possessed two additional SNPs relative to the other three; Stans, Switzerland (c.1548-1575), where two of eight genomes carried an extra SNP; and the Great Marseille Plague (1720-1722), where one of five genomes differed from the remaining four by a single SNP[48]. Importantly, as indicated in the main text, human DNA analysis previously conducted on individuals LUN369U and LUN099 suggests that they were full siblings who died at approximately 7-8 years and 15-20 of age, respectively. Hence, the chronological interval between LUN369U and LUN099 was probably no greater than 20 years, corresponding to the approximate maximum reproductive lifespan of late medieval women, given that both individuals were born to the same mother. To play on a safe side, we may extend the approximate chronological upper boundary of LUN099 to c.1430, thus, dating it to c.1413-c.1430 and potentially associating it with one of the following outbreaks: 1413-1414, 1421-,1422 and 1430-1431[54].

**Lund *LUN371* genome**

LUN371 genome is positioned on the same internal sub-branch as the previously published genome from Starnberg (Bavaria) STA001[60], associated with Keller *et al*. 2026 with one of the following outbreaks: c.1420, 1430, 1439. LUN371 sits on a terminal branch 3 SNPs derived from STA001. It could, again, imply that both STA001 and LUN371 are associated with the same wave. In this case, LUN371 could, in theory, be associated with one of the following (discussed in the previous section): 1421-1422, 1430, 1439-40. If, however, LUN371 is associated with a later outbreak than STA001, then we may extend the upper chronological boundary to 1451-1452.

Hence, there is an overlap in chronological approximation between LUN112/099 on the one hand and LUN371 on the other. Because none of these genomes comes from a precisely dated burial, it is impossible to associate the respective genomes with particular outbreaks.

**Lund *LUN361U* genome**

LUN361U is positioned in a clade with the previously published genomes X52, G371 and A146x3011 (all Denmark) and the low-coverage genome G701 excluded from analyses in this study[48,73]. LUN361 is a placed low-coverage genome, and hence its positioning is approximate. We may still use about 7 SNPs separating it from G701/X52/G371 as a rough guideline, and hence estimate that it could be about 30 years younger than these, thus hypothetically place it within the chronological range of c.1483-c.1495. If this is true, then LUN361 could potentially be associated with one of the two following outbreaks documented in Sweden in that period: 1484-1485 and 1495[54]. However, because this is a placed genome, we may, to play on a safer side, extend the chronological boundary backwards and include 1451-1452, (right around 1451 - the lower boundary of the G701/X52/G371 cluster by Keller *et al*. 2026), 1454-1455, 1464-1466 and 1474-1475 outbreaks, reported in Sweden, as two additional candidates to be associated with LUN361[54].

**Branch 1A3 Polytomy and 1A3 genomes Lund *LUN409*, Eindhoven EIN2927, and Trondheim *SK25*, *SK111* and *SK079***

The so-called 'Little Polytomy', birthing two major sub-branches and at least two (to our current knowledge) short-lived clades, and referred to by Keller *et al*. as Node 16 (N16), was preceded by another polytomy birthing four branches (two internal and two terminal), on which new genomes LUN409, EIN2927 (Eindhoven), SK25, SK111 and SK079 (all from Trondheim), all belonging to 1A3 branch, are placed. Keller *et al*. 2026 dated the 'Little Polytomy' N16 to c.1450-c.1500, and potentially to the narrower bracketing of c.1470-c.1500. The node giving rise to the 1A3 polytomy (N13 according to Keller *et al*. 2026) is 1 SNPs before the Little Polytomy (N16 according to Keller *et al*. 2026), and hence, it may have theoretically emerged c.1440-c.1490, or perhaps within the c.1460-c.1490 range.

LUN409 is a placed genome, positioned on a sub-branch with no additional genomes, about 7 SNPs derived from the 1A3 polytomy node. However, the SNP at pos. 1549630, leading to the Little Polytomy is not covered in LUN409. Therefore, the possibility that it represents a descendant of the Little Polytomy instead cannot be excluded. The lack of additional and indeed dated genomes makes any approximation of substitution rate on that sub-branch impossible, and this fact complicates placing LUN409 in an approximate dating range. The only hint is that the Lund Trinity Church graveyard went out of use with the Reformation of 1536, yielding 1536 as the absolute upper boundary. The presence of about 13 SNPs may place the lower boundary to the late 15th/early 16th century. Judging from its phylogenetic positioning, LUN409 is unlikely to be older than LUN361, placed on an older sub-branch, and it is likely that the two genomes are associated with different outbreaks. As suggested, LUN361 may have been associated with either the 1484-1485 or the 1495 outbreaks. Assuming that LUN409 is (1) associated with a later outbreak than LUN361 and (2) associated with a pre-1536 outbreak, we may hypothetically narrow it down to the range c.1495-c.1536. Within this bracketing, only the 1500, 1504, and 1508 outbreaks[54] may be identified as plague with a degree of certainty. The 1529-1530 outbreak, reported in Denmark, can most likely be identified as the 'English sweat', i.e. a different disease caused by a yet unknown pathogen, although it is possible that there was a concurrent plague outbreak in southern Norway[74,75]. It was not until 1548 that the next plague outbreak was documented in Sweden. Obviously, the lack of surviving/known documentation does not mean Sweden was spared of plague in the period c.1509-c.1547; all it means we are not aware of documented outbreaks in that period that LUN409 could potentially be associated with. Hence, it may be safe to conclude that LUN409 could be associated with one of the following outbreaks: 1500, 1504, 1508 and potentially an undocumented outbreak in the period c.1509-1536.

Branch 1A3 is characterized by a bifurcation into two sub-branches, the one with leading to the Eindhoven (EIN2927) genome and previously published Cambridge NMS002 (dated by Keller *et al*. 2026 to the c.1458-c.1529 range) and Riga G488 (dated by Keller *et al*. 2026 to the c.1549-c.1623 range), and another one leading to the Trondheim genomes cluster (SK025, SK111 and SK079). There are three or four SNPs between the 1A3 polytomy node and the EIN2927/NMS002/G488 - SK025/SK111/SK079 bifurcation node, meaning the latter may have emerged c.1450-c.1515, or perhaps c.1470-c.1515 (assuming the 1A3 polytomy node emerged c.1440-c.1490, or perhaps more specifically c.1460-c.1490).

SK025/SK111/SK079 share two clade-defining SNPs, which may position them in the c.1458-c.1530 range, or perhaps into the narrower bracketing of c.1478-c.1530. The approximate *ante quem* date of c.1530 is strengthened by the *ante quem* date of 1538 for Cambridge NMS002 (on the basis that the associated cemetery, Augustinian Friary, Cambridge) went out of use in the same year), occupying a more derived position than SK025/SK11 in relation to their most recent common ancestor. Plague outbreaks for Norway, in general, are reported in 1486, 1500, as well as more specifically in 1521 (South Norway), 1525 (south-eastern Norway) and 1529-1530 (South Norway)[75,76]. Although the three latter outbreaks (1521, 1525 and 1529-1530) were reported outside of the region where Trondheim is situated, it does not mean that they were confined to southern Norway, never reaching other parts of the country, including Trondheim. Hence, SK025/SK111 may in theory be associated with any of these outbreaks.

SK079, a placed low-coverage genome, appears to be 10 SNPs derived from SK025/SK111, although these SNPs are to be taken with a caveat. Assuming that at least the majority of these SNPs were indeed true, then SK079 must be associated with a later outbreak than SK025/SK111 (potentially belonging to the c.1486-c.1530 range) - perhaps some 40 years after the outbreak associated with SK025/SK111. This may push the dating of SK079 to c.1525-c.1570. The 1525 and 1529-1530 have already been listed in the previous paragraph; hereafter, we hear about the subsequent outbreaks in 1547-1548 in South Norway, and 1565-1566 all over the country, including Trondheim[75]. There is, however, another possibility that many of the 10 estimated SNPs are false and SK079 might be identical with SK025/SK11 and therefore is associated with the same outbreak.

Three further SNPs separate the EIN2927/NMS002/G488 clade from the common ancestor with SK025/SK111/SK079, thus potentially placing the latter in the chronological range of c.1470-c.1530, or perhaps more narrowly, c.1490-c.1530. EIN2927 is the first offshoot of this clade, having accumulated a certain number of SNPs, which is hard to estimate due to low coverage: it could be as many as 12, but in reality, it could also be much fewer than that. In absence of any additional information, phylogenetic, archaeological or radiocarbon, capable of shedding any further light on its context, we may place EIN2927 into the wide chronological bracket of c. 1490-c.1620.

Association of EIN2927 with a particular outbreak is compounded not by our inability to narrow down the wide chronological range, but also by the scarcity of references to plague outbreaks in that particular town. Some other urban centres in nearby regions provide such references. Thus, we hear about plague outbreaks in Tilburg (33 km away) in 1533, 1574-1576, 1578, 1585-1587, 1594, 1598-1604, 1616, 1625-1626; in, 's-Hertogenbosch (Den Bosch) (36 km away) in 1531, 1540, 1554, 1558 1574-1576, 1579, 1606-1607, 1616-1617, 1624-1626 and 1629; in Heusden (42 km away) in 1495, 1508-1509, 1518, 1558, 1603-1604, and 1624-1625; in Breda (60 km) in 1500, 1502, 1504, 1515-1516, 1519-1521, 1532, 1534-1535, 1541, 1546, 1574, 1585, 1587, 1589, 1593, 1599, 1602-1604, 1624-1625, 1627, and 1629-1630[77,78].

### Summary

With all these caveats and uncertainties in mind, the tentative associations between the new genomes and possible historical outbreaks may be schematically presented in the **Supplementary Table S8**.

**Supplementary Note 6: Phylogenetic placement score and approach**

In order to evaluate the placement of downsampled genomes into the phylogenetic tree, we developed a basic placement score based on normalised distances between nodes, which can be defined as follows:

If we denote the set of all the branch lengths between node A and B on the tree as,

$$s = \{b_1, b_2, \dots, b_n\}$$

Where each $b_i$ represents the length of a branch on the path between node A and B then we can represent the sum of all of the branch lengths (patristic distance) as the following,

$$d = \sum_{i=1}^{n} b_i$$

$d$ is the summation over all branch lengths $b_i$ in the set $s$. The maximum distance from node A to any other node on the tree (denoted by $m$) can be given by the following,

$$d_x = \sum_{i=1}^{n_x} b_{ix}$$

$$\forall d_x \in D,\ d_x \leq m$$

where $n_x$ is the number of branches on the path from node A to node x, and $b_{ix}$ is the length of the $i$-th branch on this path. Thus, the placement score based on normalised distances between nodes can be defined as,

$$p_{score} = \frac{d - d_{min}}{m - d_{min}}$$

However, given the minimum distance between node A and any other node on the tree ($d_{min}$) is 0, the placement score ($p_{score}$) can be simplified to the following,

$$p_{score} = \frac{d}{m}$$

The result is a score between 0 and 1 where 0 represents a placement that is identical to the original genomes placement in the phylogenetic tree and 1 represents the maximum possible distance from the original placement and as such, the worst possible placement in the tree.

**Validating phylogenetic placement method**

To validate the efficacy of pathPhynder[79] in placing low coverage *Y. pestis* genomes, we strategically chose four high coverage ancient genomes Ber45, LUN252, MAN008, SK25 (~46x, ~64.6x, ~25x and ~46.5x mean coverage, respectively); these genomes were selected from regions of the tree where we anticipate our low coverage genomes may fall. The fastq files for all genomes were then imported into Julia[80] using FASTX.jl[81] (v2.1.2) and CodecZlib.jl (v0.7.4). Subsequently, the total number of reads was randomly downsampled without replacement, ranging from 40% to 0.078125% of the total reads.

The downsampled fastq files underwent processing using the nf-core eager pipeline (see Processing ancient genomic data). The resulting bam files were then utilised for placement (see Phylogenetic placement). To ensure the robustness of our results, the phylogenetic placement was repeated 100 times. Each of the trees was imported into Julia with Phylo.jl[82] (v0.5.2), and a placement score (see Placement score) for each of the downsampled genomes was calculated. After evaluating the placement in all trees, a mean placement score was calculated for each downsampled genome (**Supplementary Figure 11)**.
