## Supplementary Figures for "The Genomic Landscape of Post-Black Death Epidemics in Northern Europe and the Caucasus"

**Contributed equally*

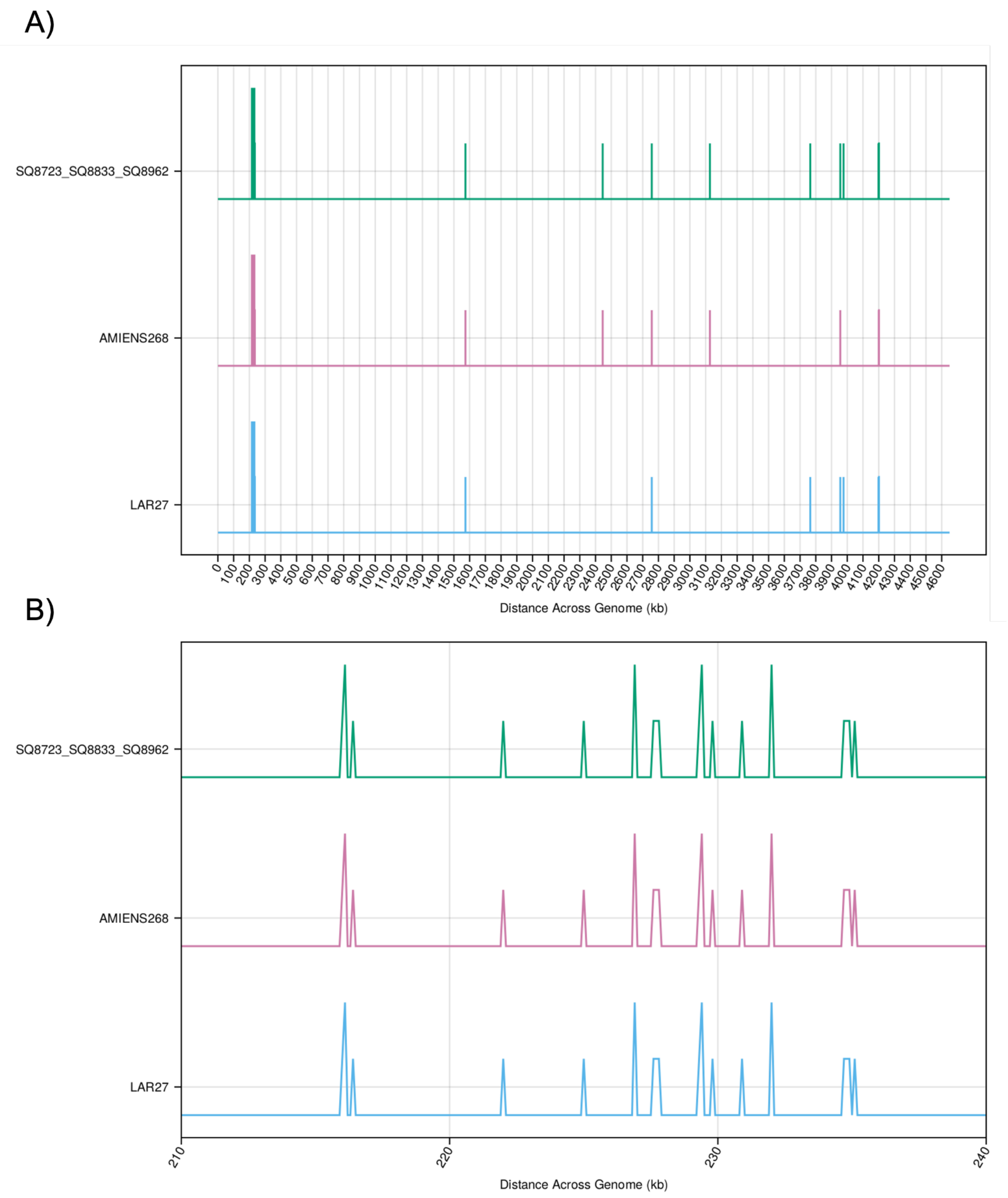


**Supplementary Figure 1:** Distribution of mutations identified in problematic genomes. A) The distribution of mutations across the genome observed for SQ8723_SQ8833_SQ8962, AMIENS268 and LAR27; all genomes that displayed unusually long terminal branch lengths for their estimated age in the *Y. pestis* phylogeny. B) Density of mutations across each of the three genomes between the coordinates 210kb and 240kb of *Y. pestis* CO92.

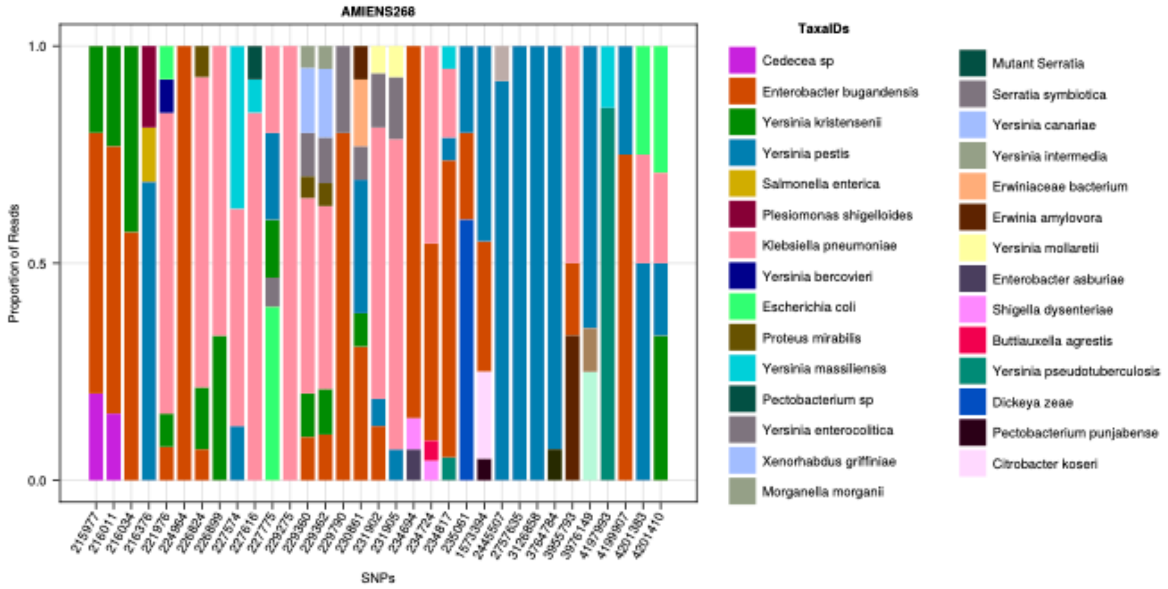


**Supplementary Figure 2:** Species represented in the top hits assigned by blastn to reads overlapping with the mutations identified in *Y. pestis* genome AMIENS268.

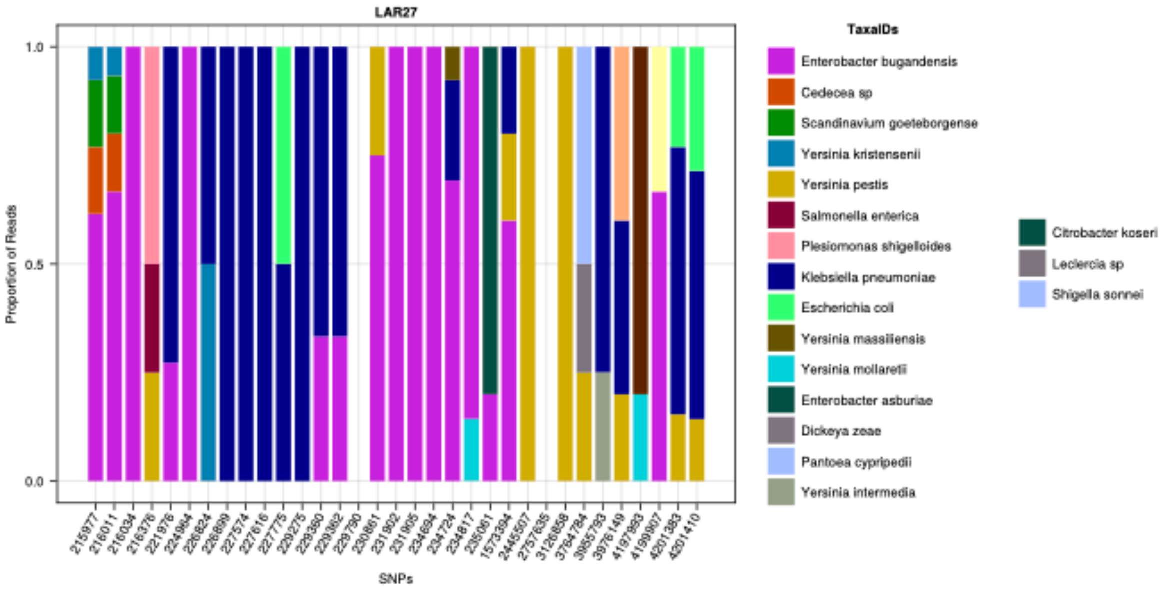


**Supplementary Figure 3:** Species represented in the top hits assigned by blastn to reads overlapping with the mutations identified in *Y. pestis* genome LAR27. White bars give reads with no taxa confidently assigned.

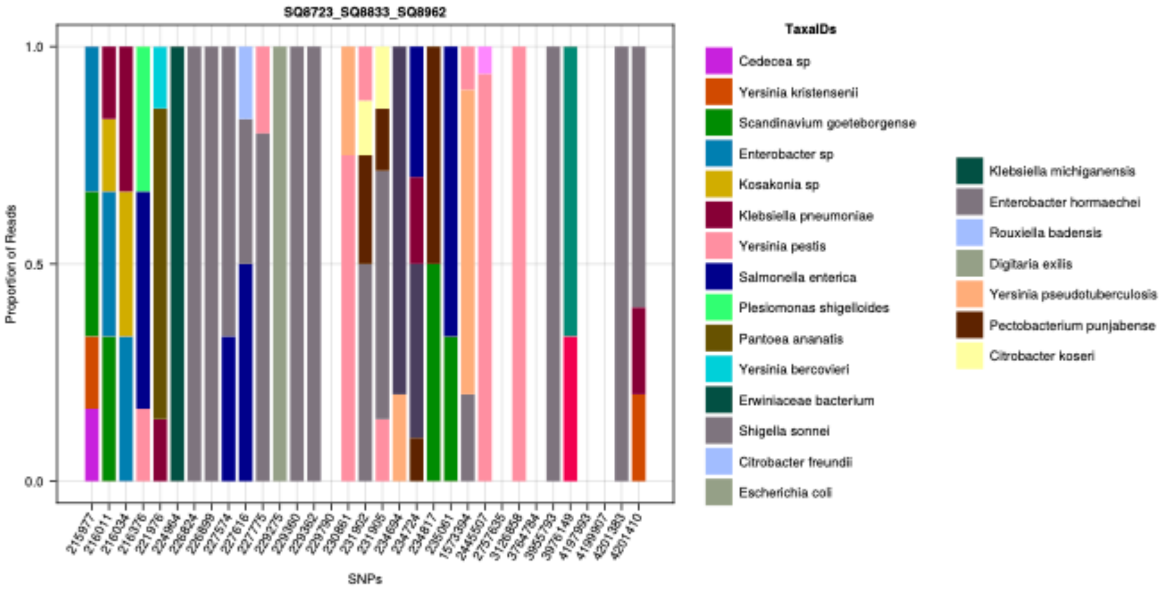


**Supplementary Figure 4:** Species represented in the top hits assigned by blastn to reads overlapping with the mutations identified in *Y. pestis* genome SQ8723_SQ8833_SQ8962. White bars give reads with no taxa confidently assigned.

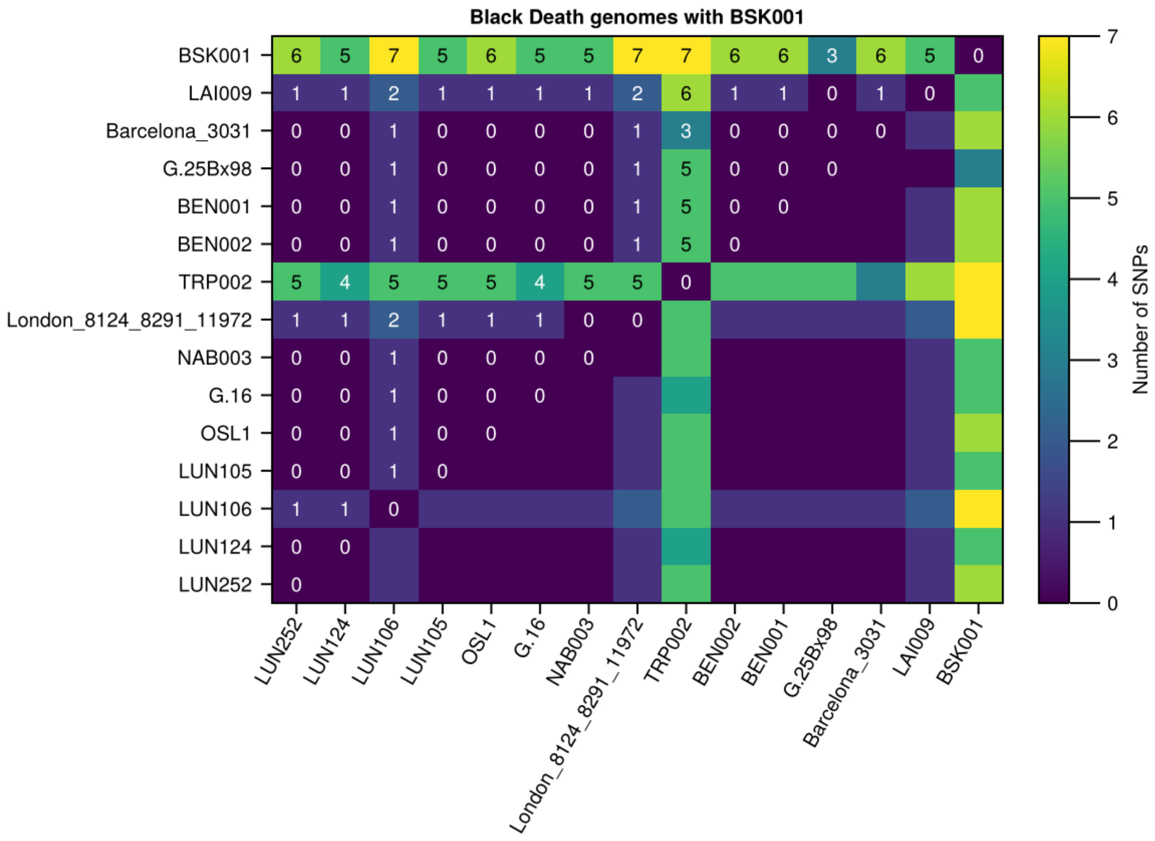


**Supplementary Figure 5:** Heatmap providing the pairwise SNP differences between *Y. pestis* genomes recovered from the Black Death, including BSK001. Differences are provided only for overlapping SNP positions, requiring the SNP to be present in both target and comparator genome.

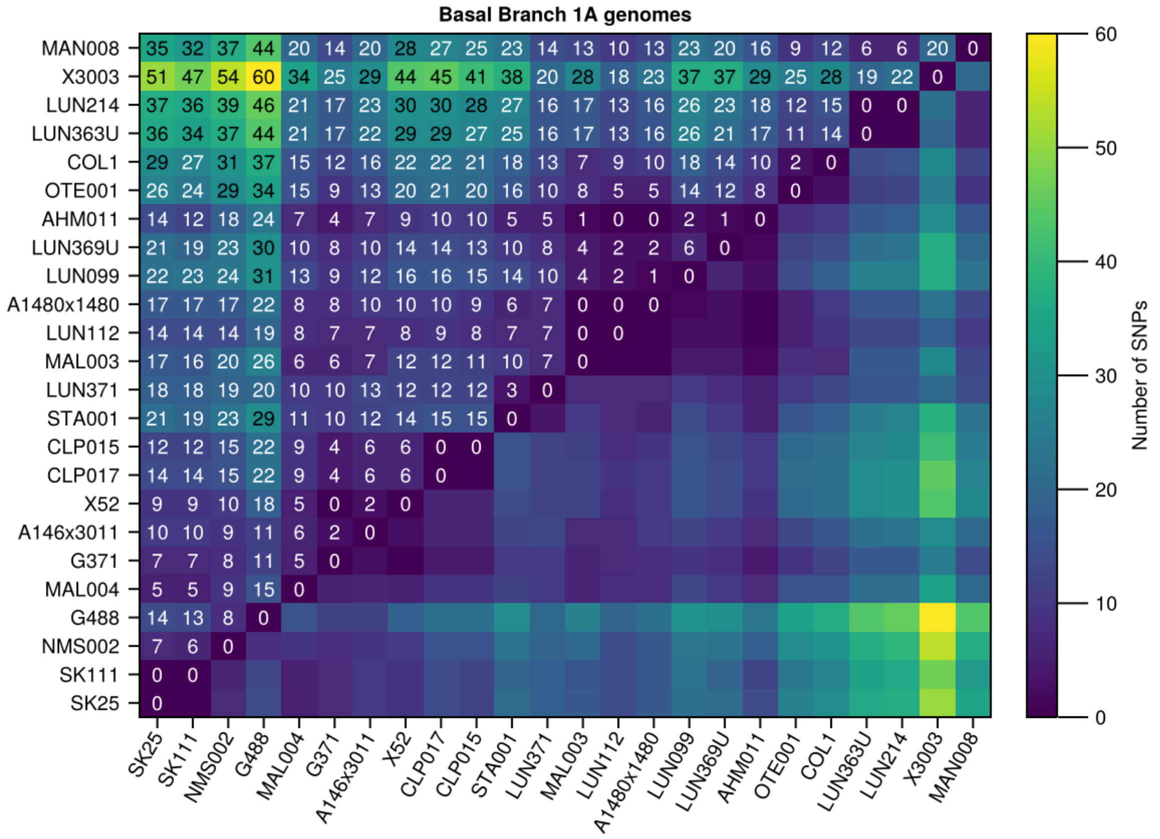


**Supplementary Figure 6:** Heatmap providing the pairwise SNP differences between *Y. pestis* genomes from basal branch 1A. Differences are provided only for overlapping SNP positions, requiring the SNP to be present in both target and comparator genome.

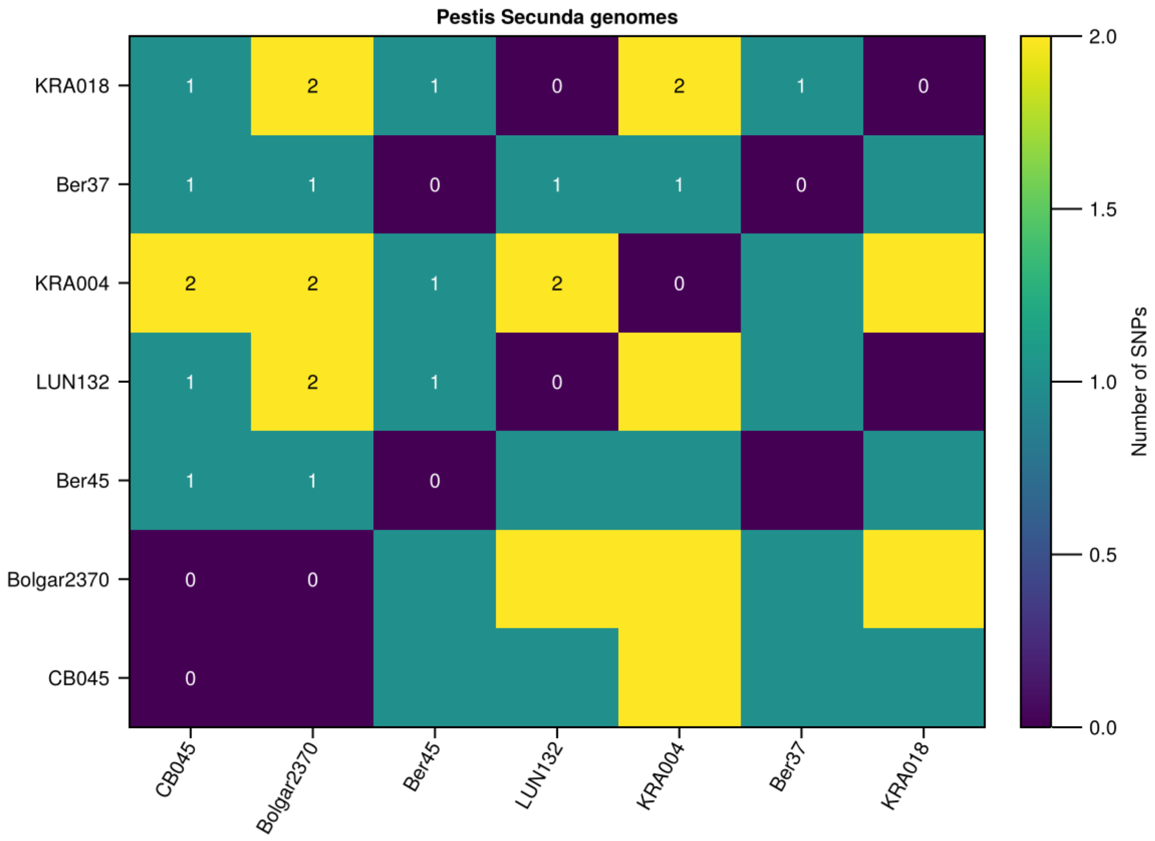


**Supplementary Figure 7:** Heatmap providing the pairwise SNP differences between *Y. pestis* genomes recovered from the *pestis secunda*. Differences are provided only for overlapping SNP positions, requiring the SNP to be present in both target and comparator genome.

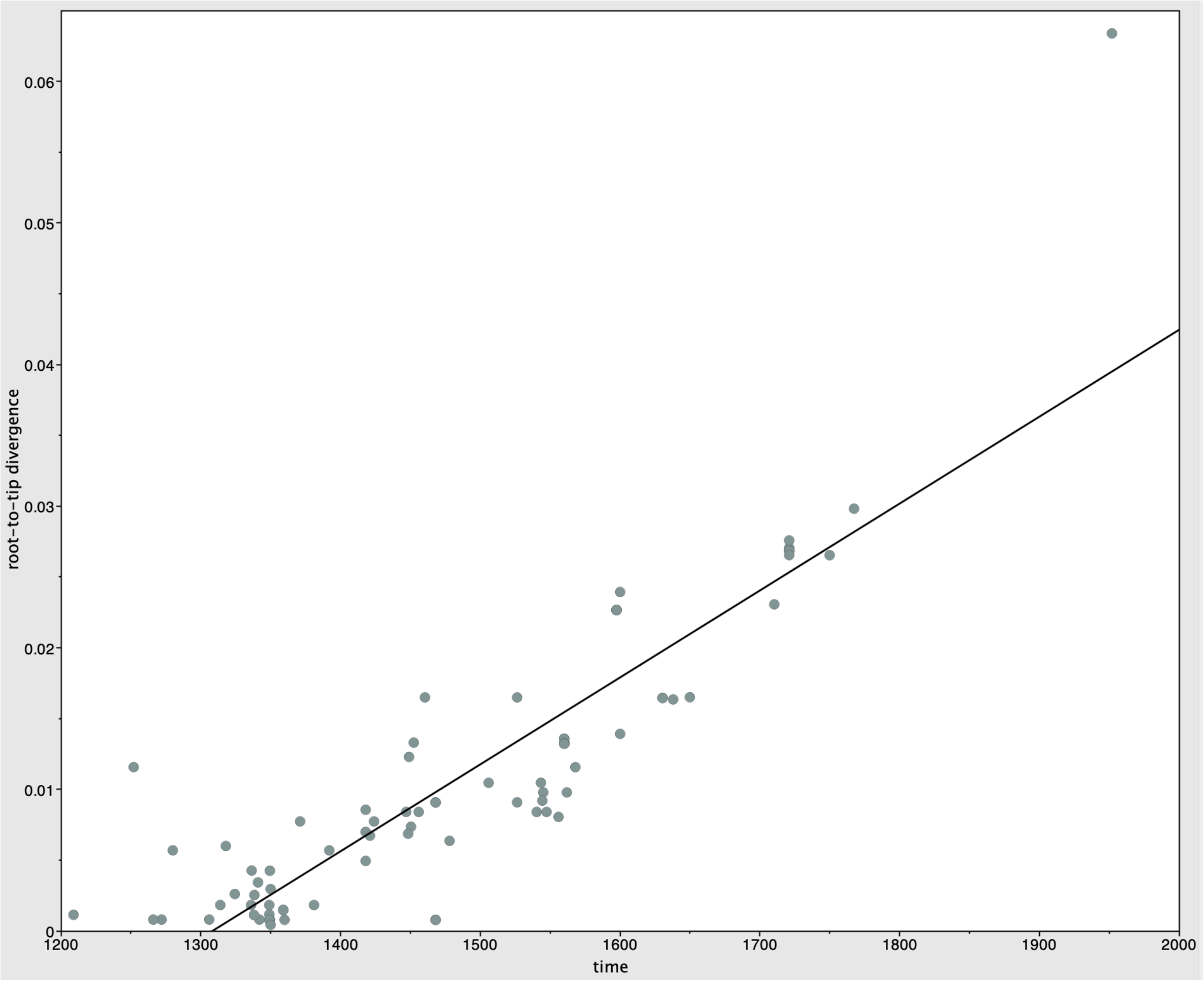


**Supplementary Figure 8:** Root to tip temporal regression over Second Plague Pandemic genomes generated with TempEst. A plot depicting the root to tip correlation for only the Second Plague Pandemic genomes of the Black Death, the *pestis secunda* and Branch 1A of the tree, R^2 = 0.7791. The BactDating date randomisation test yielded a p-value of $p<1.00e^{-4}$.

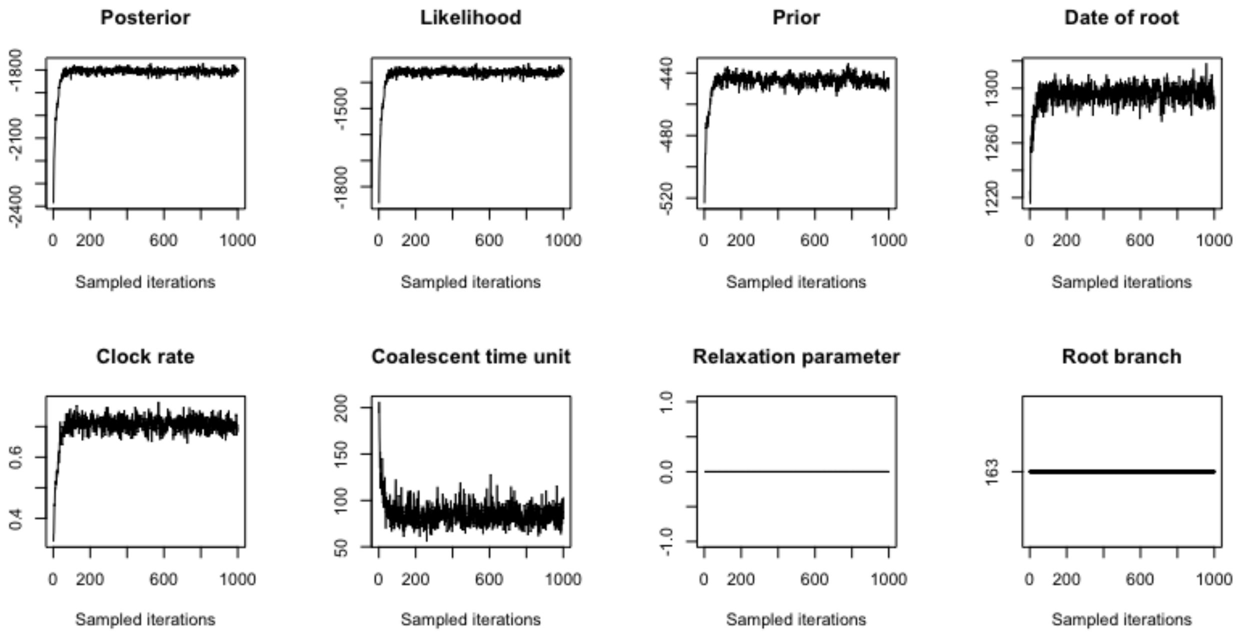


**Supplementary Figure 9:** BactDating MCMC trace plot demonstrating full convergence of the analysis.

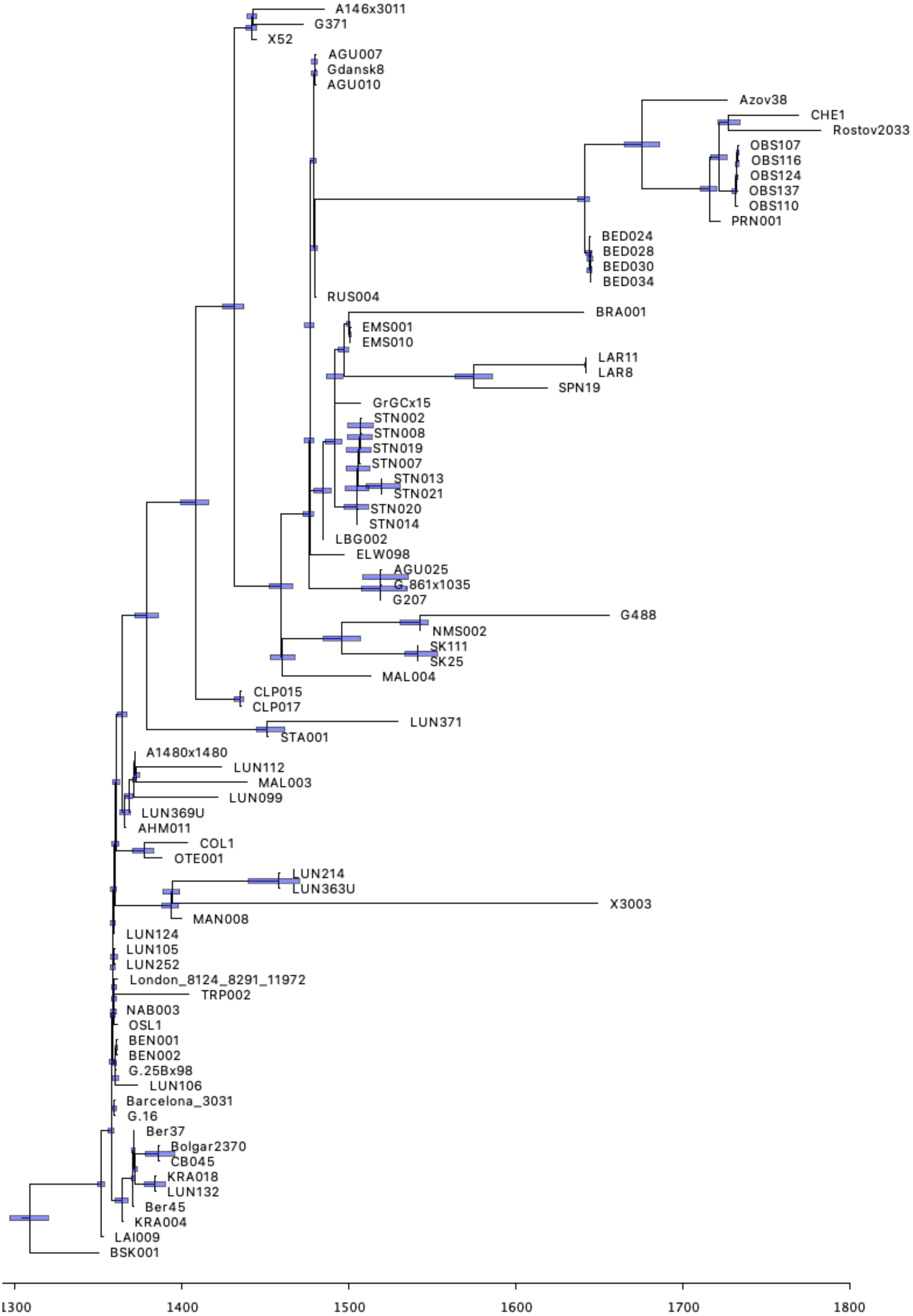


**Supplementary Figure 10:** BactDating inferred time tree under a strict gamma model. Blue bars indicating 95%CI for ancestral dates.

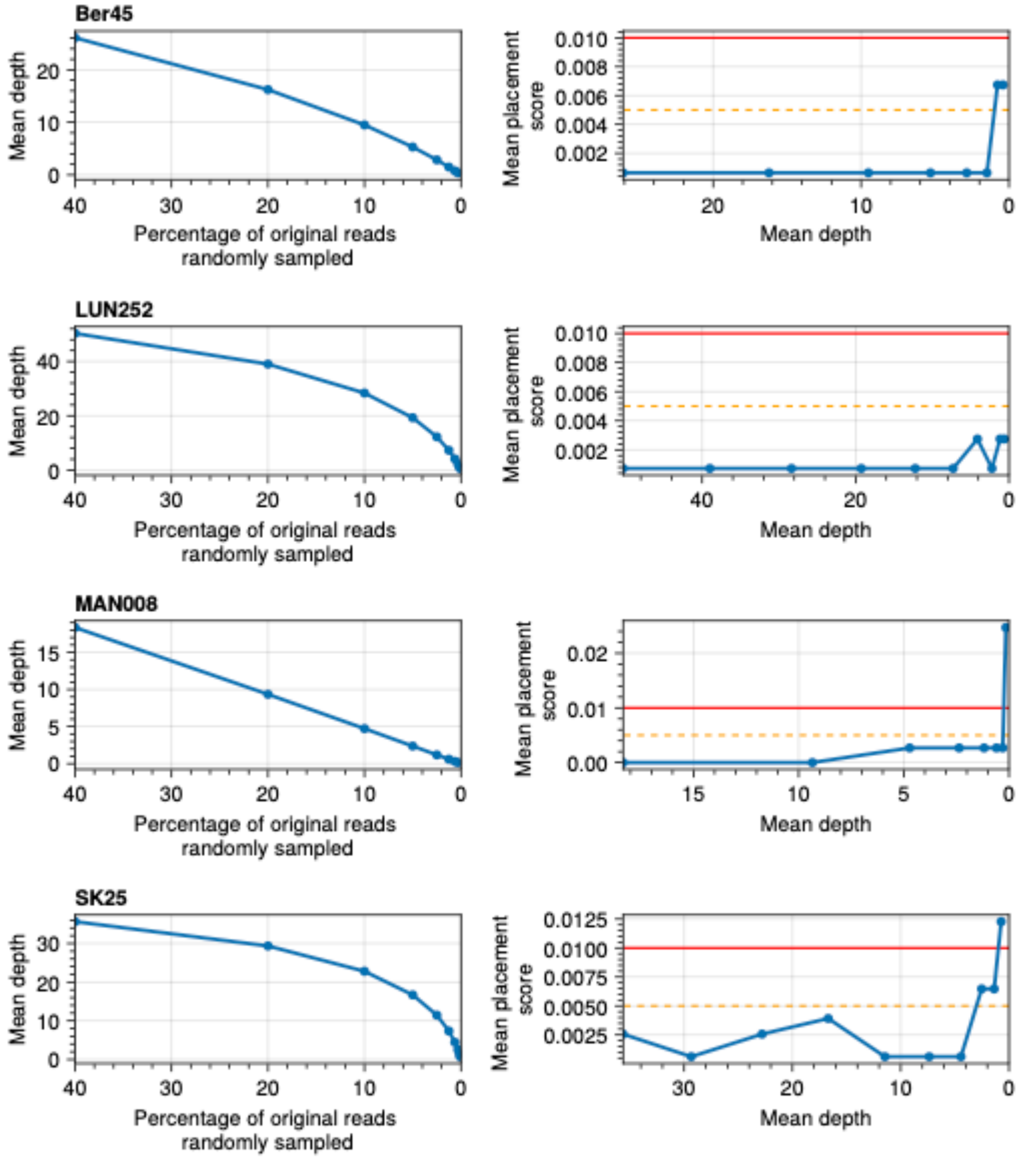


**Supplementary Figure 11:** Validation of the phylogenetic placement approach via coverage down-sampling of high quality ancient genomes. Each row of the multi-panel figure provides the approach tested on an example genome as given in the header. Panels on the left show the mean coverage of the sample after random downsampling and the panels on the right show the mean corresponding score after being placed in the tree. Below the orange line indicates a placement almost identical to the original, between the orange and red lines represents a good placement score, while above the red line indicates a placement quite distant to the original, true placement.

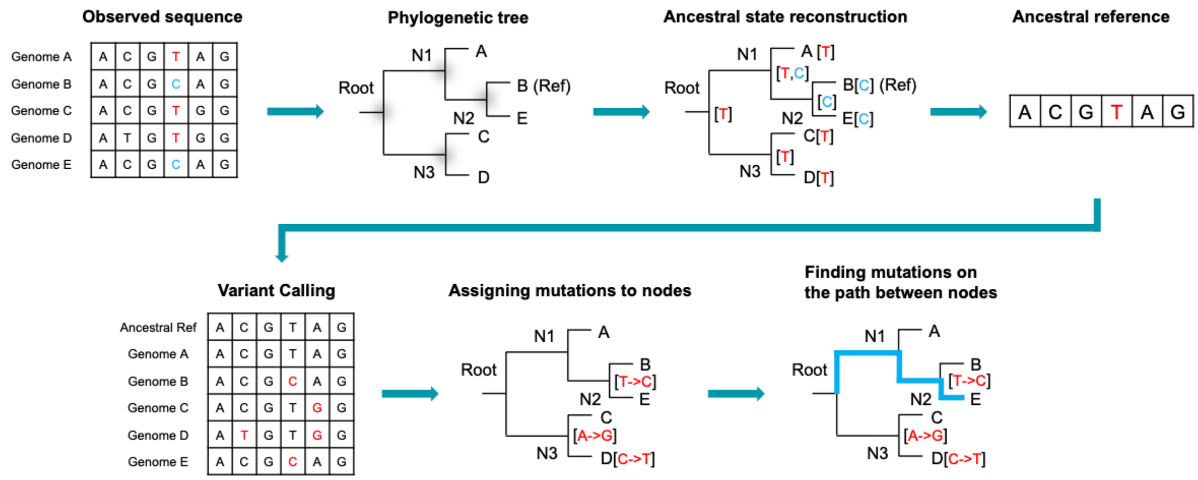


**Supplementary Figure 12:** Schematic documenting the workflow for our Ancestral Reference Mapping (ARM) approach (see **Methods**).

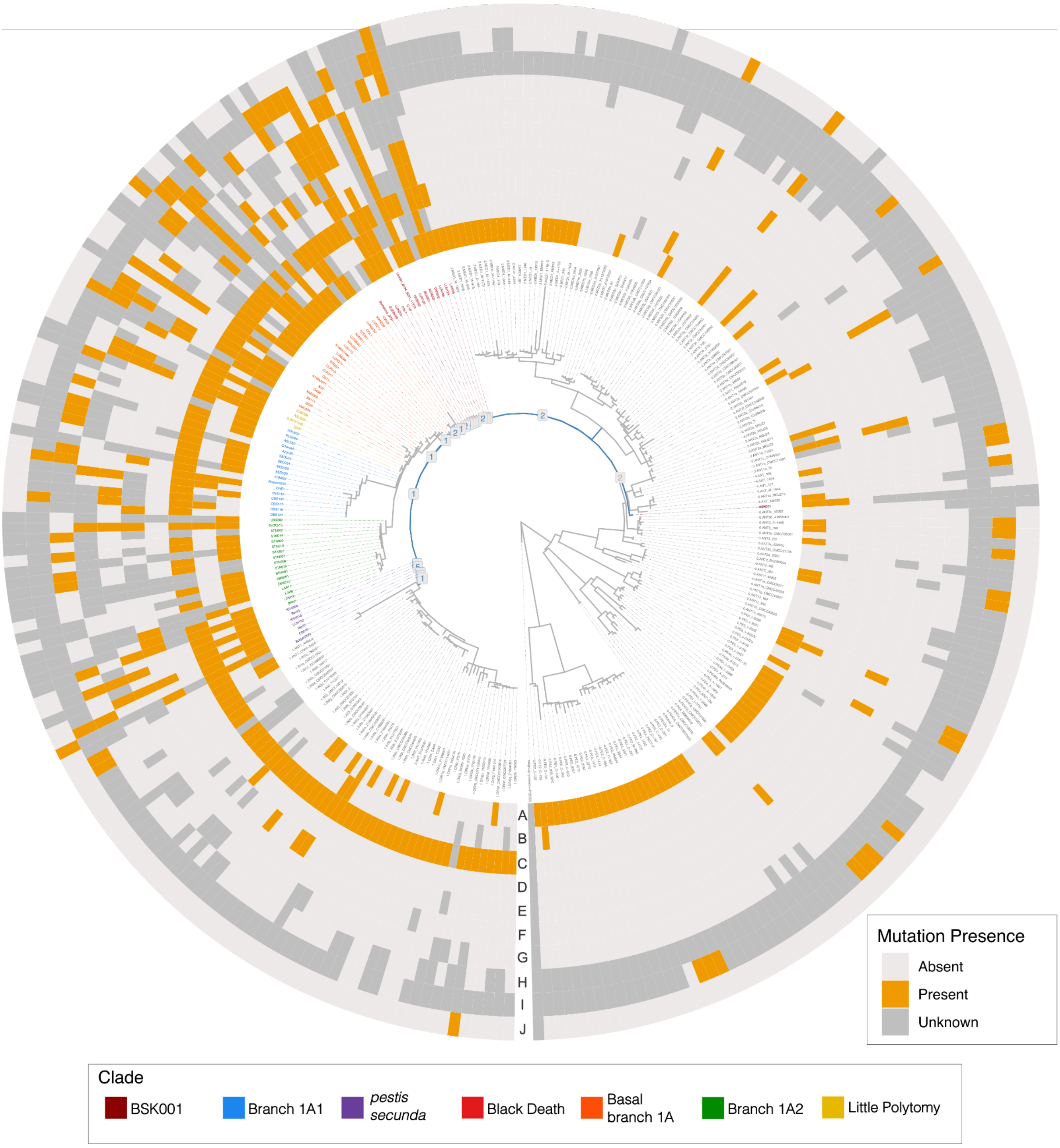


**Supplementary Figure 13:** Radial maximum likelihood *Y. pestis* phylogeny and heatmap depicting the presence and absence of intergenic mutations and indels relative to an inferred ancestral reference. The path from the Great Polytomy node to the most basal *pestis secunda* node is shown in blue with the number of mutations assigned to each node shown as labels. Coloured tip labels indicate which phase of the pandemic samples belong to. Rings of the heatmap correspond to: A) the presence and absence of the +1A insertion at 1234971; B) the presence and absence of the +1A insertion at 2916749; C) the presence and absence of the A->G mutation at 1871476; D) the presence and absence of the +1T insertion at 2639544; E) the presence and absence of the +1C insertion at 3459494; F) the presence and absence of the +1A insertion at 2764536; G) the presence and absence of the +1A insertion at 2799914; H) the presence and absence of the -1G deletion at 4000717; I) the presence and absence of the A->G mutation at 3081606; J) the presence and absence of the +2TT insertion at 2415025.

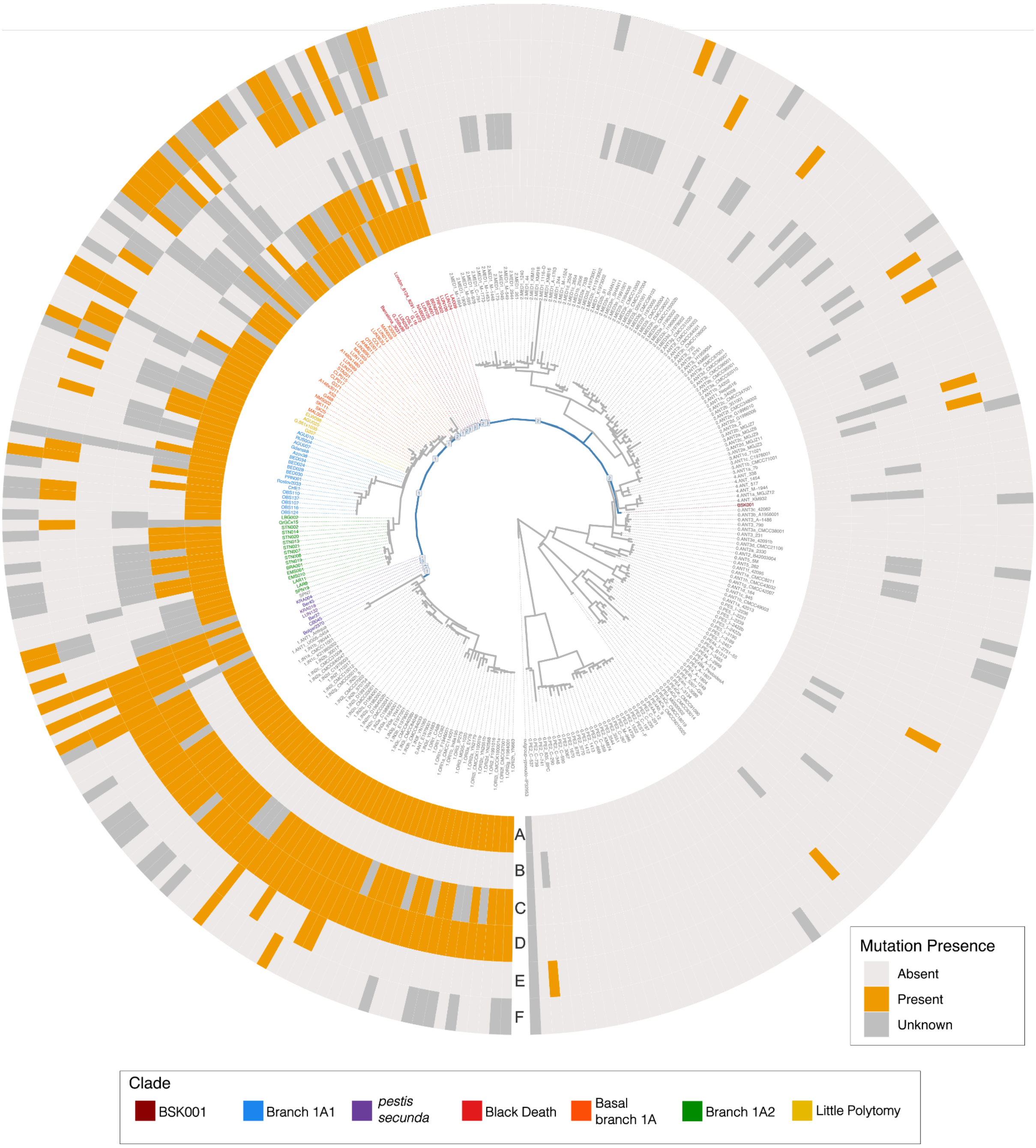


**Supplementary Figure 14:** Radial maximum likelihood *Y. pestis* phylogeny and heatmap depicting the presence and absence of nonsynonymous mutations and indels relative to an inferred ancestral reference. The path from the Great Polytomy node to the most basal *pestis secunda* node is shown in blue with the number of mutations assigned to each node shown as labels. Coloured tip labels indicate which phase of the pandemic samples belong to. Rings of the heatmap correspond to: A) the presence and absence of the T->C mutation in the YPO_RS01855 gene; B) the presence and absence of the -1A deletion in YPO_RS10230 gene; C) the presence and absence of the T->G mutation in the *hmsR* gene; D) the presence and absence of the G->T mutation in the YPO_RS10940 gene; E) the presence and absence of the +1T insertion in the YPO_RS17990 gene and F) the presence and absence of the +1T insertion in the YPO_RS19400 gene.
